# Benthic biodiversity hotspots in the Weddell Sea: Bridging geomorphology, biogeography, and oceanography across scales

**DOI:** 10.64898/2026.04.12.717989

**Authors:** Miao Fan, Autun Purser, Bingbing Wei, Tea Isler, Natalie Cornish, Boris Dorschel, Matthias Wietz

## Abstract

Benthic ecosystems are shaped by seafloor structure, yet linking geomorphology and biology across environmental gradients remains challenging. Here, we integrate seafloor imagery, multiscale bathymetry, and predictive modelling to quantify benthic biodiversity and its environmental drivers along the Powell Basin flank of the Antarctic Peninsula. Steep geomorphological landforms (terraces and steep slopes) hosted maximal densities and distinct communities, with significant enrichment of corals, sponges, ophiuroids, and sea pens. The marked congruence of slope across bathymetric resolutions enabled regional upscaling, estimating a standing stock of ∼96 billion individuals across 7,400 km² of the basin flank. Decadal oceanographic models indicate that benthic densities peak in cold bottom water below −0.15 °C. We identified four biodiversity hotspots (∼33 km^2^) with slopes >35° and depths >1,700 m, with up to threefold higher densities. Hotspots align with the pathway of Weddell Sea Deep Water, where thermal decoupling between bottom and overlying waters indicates dynamic hydrography along steep, biodiverse terrain. Furthermore, contrasting sea-ice cover and stratification regimes suggest distinct cryo-pelagic-benthic coupling around hotspots. The concentration of biodiversity through seafloor geomorphology and ocean circulation bridges Antarctic benthic ecology from habitat to biogeography, with implications for monitoring change and guiding conservation in the warming Weddell Sea.

## INTRODUCTION

The Antarctic Weddell Sea, a key region for global ocean circulation and climate, supports dense and diverse communities along its continental slope^1,2^. This biological wealth is embedded in a unique oceanographic setting: the Weddell Sea is the dominant source of Antarctic Bottom Water, controlling deep-ocean stratification and ventilation, and acting as a major carbon sink^3^. However, this system faces a triple threat from climate change: attenuation of bottom currents, altered primary production, and modified bottom water export^4–6^. These processes are projected to affect benthic communities by altering the physical and trophic setting in which they are embedded.

Understanding how benthic communities align with these dynamics requires a detailed assessment of their foundational habitat. Across the global ocean, seafloor geomorphology is an important driver of benthic life, modulating near-bottom hydrodynamics and organic matter supply^7^. Structurally complex terrain – such as slopes, canyons, and ridges – often supports greater biomass and species richness than flat, homogeneous areas^8–11^. Complex seafloor and hard substrates with high slope and roughness can enhance current speeds and food availability, favoring suspension feeders such as corals and sponges^12–14^. These geomorphological controls align with broader depth, geographic, and hydrographic gradients^15–17^.

Benthic habitat mapping is a key tool to resolve these relationships, integrating geomorphology, biology, and oceanography^18–21^. This includes the delineation of seafloor landforms; spatial units that synthesize topographic features into habitat types^22–27^. The mapping of seafloor landforms advances the quantitative understanding of benthic habitats, especially in combination with ecological modelling^28–31^. Consistent with these global patterns, Antarctic benthic communities are strongly shaped by seafloor topography, with elevated biomass and diversity on steep, complex terrain^32–36^. This fine-scale habitat heterogeneity links local ecology to broader patterns of Antarctic biogeography^37–39^. The diverse benthic communities on the shelves and slopes around the Antarctic Peninsula provide key ecosystem services but are threatened by habitat loss^40–42^. However, fine-scale terrain controls on Antarctic benthic communities remain poorly resolved, as seafloor landforms have only been mapped at coarser resolution^43^.

A key challenge is bridging the scales: how do local geomorphological controls on biodiversity translate to the frameworks relevant for biogeography and conservation? Underwater vehicles like the Ocean Floor Observation and Bathymetry System (OFOBS) leverage centimeter-scale bathymetry and imagery to document benthic ecosystems in the deep sea^44–46^. However, their limited spatial coverage constrains broader biogeographic inferences. By contrast, decameter-scale ship bathymetry captures only coarser terrain features and may miss finer gradients^47,48^. Linking benthic biology and geomorphology across scales, including the predictive upscaling of local empirical data, can promote biodiversity research, habitat mapping, and ecosystem management^49–51^. This includes identifying biodiversity hotspots and their environmental setting^30^ – habitats with diverse ecosystem services that could be prioritized for conservation.

In this study, we integrated bathymetric, imaging, and oceanographic data from the Powell Basin of the Antarctic Peninsula to (i) characterize seafloor landforms in high resolution, (ii) assess their benthic life and environmental connections, and (iii) establish a larger bio-oceanographic framework through biogeographic modelling. In the Powell Basin, a key region for Antarctic Bottom Water formation and export, complex seafloor topography and strong currents sustain diverse benthic communities^52–55^. The region is experiencing pronounced climate-driven change while detailed baseline data are scarce, limiting the prediction and mitigation of ecological impacts. By quantifying 10 key benthic taxa across seven landforms, we reveal local habitat-biology relationships, upscale those along the continental slope, and identify benthic hotspots along major Antarctic water masses. The evidence that seafloor structure and oceanography jointly shape benthic biodiversity resolves key dynamics and drivers of the Antarctic benthos, supporting future ecosystem mapping and conservation^56–58^.

## METHODS

### Study area and design

On expedition PS118 of RV Polarstern in 2019^59^, benthic imagery and bathymetry were collected using the Ocean Floor Observing and Bathymetry System (OFOBS) on all three dives (designated 39-1, 69-1, 81-1) on the northern Powell Basin flank, covering complex topography between 500 and 2200 m depth (Figure 1, Supplementary Table S1). A forward-looking sonar allowed operating over the very steep terrain, generating a unique dataset along the flank walls. We then classified seafloor landforms from high-resolution bathymetry to characterize the geomorphological setting. The dives were considered as replicates of recurring landform classes, while accounting for underlying spatial gradients (depth, latitude, longitude). We then annotated and quantified benthic fauna in their geomorphic setting, followed by spatial modelling of taxon densities and the definition of biodiversity hotspots. Finally, biogeographic estimates were integrated with hydrographic models to establish a broader oceanographic context.

**Figure 1.**
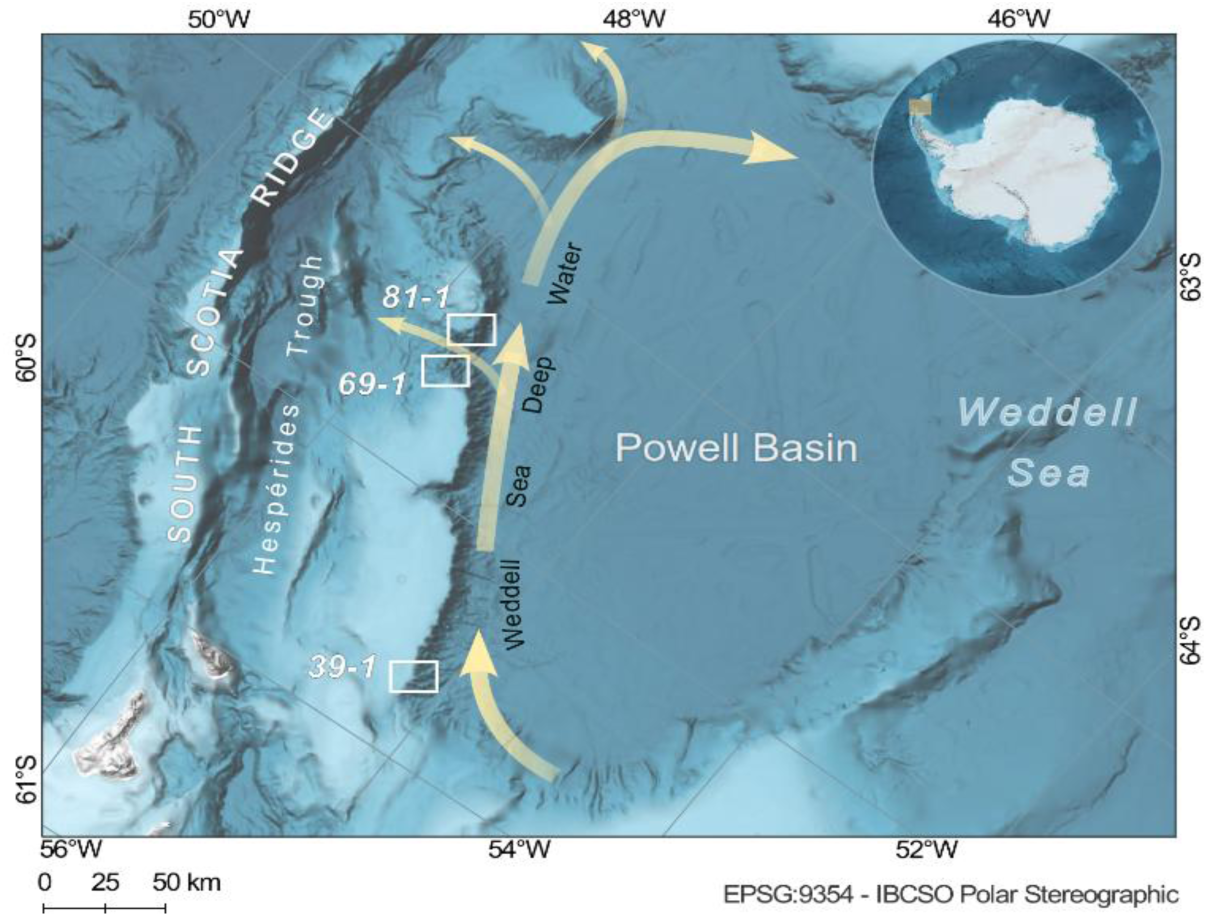
Study area and water mass pathways. The white boxes indicate the three OFOBS dives on the Powell Basin flank. The yellow arrows indicate the pathway of Weddell Sea Deep Water along the continental slope, derived from^55,60^. The basemap is IBCSO v2.0 bathymetry^61^.

### OFOBS bathymetry and imagery

The OFOBS was towed ∼2 m above the seafloor, acquiring multibeam and sidescan sonar data via an interferometric EdgeTech 2205 multiphase echosounder^50,51^. Simultaneously, a Canon EOS 5D Mark III captured 26-megapixel images (typically covering ∼4 m² of the seafloor) every ∼20s. An integrated Posidonia transponder for ultra-short baseline triangulation allowed for georeferencing with a horizontal positioning accuracy of ∼0.2% of the slant range. Each image displays three laser dots forming an equilateral triangle (50 cm) near the center, providing a spatial reference. From 7,194 photographs, 2,252 were excluded due to Posidonia failure or image quality (black frames from ascent/descent or flash malfunction), leaving 4,942 for analysis.

### Ship bathymetry

Multibeam data were acquired with a hull-mounted Teledyne Atlas HYDROSWEEP DS3 echosounder. Teledyne CARIS HIPS and SIPS v.11.4 was used to remove erroneous measurements, and to correct for sound velocity changes and tides by referencing to mean sea level. From the cleaned data, a Digital Elevation Model (DEM) was calculated at 25 m resolution. Sidescan sonar data was processed by applying altitude correction, slant range correction, bottom tracking, and automatic gain control; then mosaicked with 2 cm resolution.

### Terrain attributes and seafloor landforms

We extracted five geomorphological terrain attributes (slope, aspect, roughness, plan curvature, profile curvature) that represent standard descriptors of seafloor topography and texture^62,63^. Attributes were extracted at all available resolutions: 10 cm, 20 cm, 1 m, 2 m (OFOBS) and 25 m (ship) using a 3×3 moving-window analysis. In addition, we calculated the Bathymetric Position Index (BPI), which quantifies the elevation difference between a georeferenced cell and the average of a defined concentric reference area. Using a focal function within an annulus shape defined by inner / outer radius values, the BPI identifies morphological highs (positive values) and lows (negative values) compared to the reference. Both fine-scale BPI (inner / outer radius: 1 / 3 cells; 20×60 cm) and broad-scale BPI (inner / outer radius: 50 / 200 cells; 10×40 m) were calculated to capture local and larger geomorphic structures, respectively. Supplementary Figure S1 depicts all terrain attributes for an exemplary section of the study area.

We classified seafloor landforms from OFOBS DEMs (20 cm resolution) using the Benthic Terrain Modeler (BTM) extension for ArcGIS. Following an established classification scheme^64^ with defined BPI and slope thresholds (Supplementary Table S2; Supplementary Figure S2), we first delineated geomorphic landform zones and subsequently seven landforms: terraces, steep slopes, ridges, broad slopes, flat areas, depressions, and sand ripples. Terraces and sand ripples are compound landforms resolved by BPI, slope, and characteristic spatial patterns in high-resolution bathymetry and backscatter mosaics, as confirmed using seafloor imagery (step-like benches with local, narrow ridges and depressions on steep slopes for terraces; regular ridge–trough morphology for sand ripples). The classification was confirmed through a representative set of 371 images: all 196 taxonomically annotated images (see below) and 175 non-annotated images selected using *Random extract within subsets* in QGIS, ensuring proportional representation across landforms and dives. This sample size corresponds to a statistically representative subset (∼95% confidence, ±5% margin of error). Each image was overlaid with the BTM-derived landform map, yielding a classification accuracy of 92% (Supplementary Table S3). This framework establishes a quantitative link between geomorphological and biological patterns.

We then applied pairwise Kruskal-Wallis and Dunn’s tests with Benjamini-Hochberg correction to identify which terrain attributes differ most between landforms. The fraction of significant contrasts (i.e. all pairwise comparisons with adjusted *p* < 0.05) identified slope as the strongest discriminator (Supplementary Table S4). While the highest fraction occurred at 2 m resolution (0.62), the median adjusted *p*-value at 20 cm was three orders of magnitude lower (9 × 10⁻⁷), confirming that 20-cm resolution best delineates landforms. Roughness exhibited similar fractions (0.52–0.57), whereas aspect, plan curvature, and profile curvature (≤0.24) provided little separation. Terrain attributes at 20 cm resolution, together with depth, latitude and longitude, were then analyzed by Principal Component Analysis (PCA) to determine how landforms align with environmental gradients.

### Benthic fauna annotation and taxonomy

For a representative assessment of benthic fauna, we selected 196 images that met three criteria: (i) high quality and sharpness, (ii) optimal illumination, and (iii) uniform spacing along each dive and balanced counts per landform. Benthic organisms (larger than 2 cm) were identified to the lowest possible taxonomic level and assigned to 10 taxon groups in BIIGLE 2.0^65^ following established identification catalogues^66–72^ and expert guidance (see Acknowledgements). These taxa are common components of Antarctic benthic communities^73^ and can be reliably identified from seafloor imagery: corals, sea pens, anemones (phylum Cnidaria), ophiuroids, asteroids, crinoids, echinoids (Echinodermata), demosponges and glass sponges (Porifera), and bryozoans (Bryozoa). The higher-level grouping ensured sufficient replication for statistical analyses. Corals included stylasterid lace corals, solitary cup corals, branching octocorals, and gorgonians (for example *Thouarella*, *Primnoella*, *Primnoisis*, *Callogorgia*). Bryozoans (for example *Smittina*, *Cellarinella*, *Reteporella*) were quantified when colonies were visually distinguishable. Branching geometry and the presence of polyps supported the visual distinction of bryozoans and corals. Sponges were distinguished into demosponges (including *Stylocordyla*, *Cinachyra*, polymastiid forms) and glass sponges (including *Rossella antarctica*, *Asconema setubalense*, *Aphrocallistes vastus*) by opaque and massive versus cup- or vase-like morphology, respectively. Echinoderms included *Astrochlamys* and *Amphiura* brittle stars, *Odontaster* sea stars, *Promachocrinus kerguelensis* feather stars, and pencil urchins (Supplementary Table S5). Counts were normalized and scaled to 1 m², considering the usable seabed area for each frame using a rectification factor (accounting for altitude and perspective). Overall, our approach ensures reproducibility and interpretability while acknowledging the limitations of image-based taxonomy.

### Density and composition of benthic fauna

Taxon densities between landforms were compared by Kruskal–Wallis and Dunn’s tests in combination with Generalized Additive Models (GAMs) to capture non-linear responses to terrain gradients. Because slope and roughness strongly correlate (ρ = 0.9), we first tested whether both attributes are needed to assess densities. GAMs with slope alone, roughness alone, and both demonstrated that roughness did not improve model performance (ΔAIC ≥ −2) for any of the 10 quantified taxa. Consequently, roughness was excluded from subsequent analyses. Variance inflation factors (VIF) below 5 confirmed that all remaining variables did not exhibit critical collinearity. Densities were log-transformed, as this provided a more balanced distribution than square-root transformation (Supplementary Figure S3). A GAM including landform categories, terrain attributes, depth, latitude, and longitude was superior to a reduced model excluding landform categories (AIC 4022 vs. 4080). To test compositional variability between landforms, we ran redundancy analysis (RDA) on log-transformed densities constrained by terrain attributes, depth, latitude, and longitude. ‘Flat area’ was used as reference category because its high replication (*n* = 30) and low morphological complexity provide a neutral baseline for comparison with other landforms. Distance-based redundancy analysis (dbRDA) on Bray–Curtis dissimilarities was run for comparison, explaining 24% of community variation compared to 37% for standard RDA. To assess the individual contributions of geomorphology, depth, and site, we summarized each group by PCA and used the first two principal components per group for variance partitioning.

### Biogeographic modelling

We first compared terrain attributes derived from OFOBS and ship bathymetry via pairwise Spearman correlations, which assess consistency in relative gradients independent of absolute scale differences, identifying which attributes are robust across resolutions. Area-corrected densities of all taxa were then modelled using a negative-binomial model (function *manyglm* from package *mvabund*); once with OFOBS-derived and once with ship-derived terrain attributes alongside depth, latitude, and longitude (all z-score scaled). Negative-binomial models were used to account for overdispersed counts typical of patchy benthic communities. For taxa with directionally consistent responses, densities along the cruise track were then estimated using GAMs (package *mgcv*) with ship-derived slope and depth as predictors, allowing the spatial prediction of non-linear density–habitat relationships. Predictive performance was assessed using train–test splits comparing dives 39-1 and 69-1 with 81-1, yielding moderate correlations (r ≈ 0.31), low mean absolute error (MAE ≈ 4.2), and stable root mean square error (RMSE ≈ 7.7), consistent with the spatial patchiness of benthic fauna and other modelling studies^74,75^. Total standing stocks across the study area (≈ 12 million grid cells of 25×25 m; 7,400 km²) were calculated from the raw predictions (expressed as densities per m²) by multiplying with grid-cell area (25×25 m = 625 m²). Model uncertainty was quantified using the approximate covariance matrix of the penalized likelihood estimates, with prediction variance propagated to the response scale using the delta method.

### Biodiversity hotspots

We identified biological peaks through a hierarchical clustering and filtering approach – from individual grid cells over high-density clusters to spatially coherent hotspots. First, individual grid cells along the cruise track were spatially aggregated (500×500 m). Aggregations in which more than half of grid cells were within the top 10% of predicted density (evaluated both globally and within 500-m depth bins) were retained, and their strength defined as the fraction of “hot” cells multiplied by the standardized mean predicted density. Retained aggregations were then grouped into high-density clusters using a two-stage DBSCAN approach (eps = 1500, minPts = 4 followed by eps = 500, minPts = 3), and reassigned to the cruise coordinates using nearest-neighbor mapping (k = 1). High-density clusters within the top 10% of strength (global > 0.84, depth > 0.88) were grouped into hotspots using DBSCAN (minPts = 6000). Environmental conditions and predicted densities were compared between hotspots, high-density clusters, and non-hotspots by Dunn’s tests with Benjamini–Hochberg correction on each 5,000 grid cells (random subsampling with fixed seed).

### Hydrographic context

We extracted hydrographic variables from the CMEMS Global Ocean Physics (GLOBAL_MULTIYEAR_PHY_001_030) and Global Ocean Colour (OCEANCOLOUR_GLO_BGC_L4_MY_009_104) model products, providing 1/12° horizontal resolution at minimal bias compared to *in situ* observations; well representing currents and water masses^76^. Potential temperature, potential bottom temperature, velocity components, mixed-layer depth, sea-ice concentration, and chlorophyll-a were downloaded in monthly resolution between 01-01-1993 and 01-04-2019; corresponding to the full temporal CMEMS coverage until the expedition. Cruise coordinates were three-dimensionally matched to CMEMS data using nearest-neighbor mapping (k = 3), only considering matches with a maximum horizontal distance of 10 km and a maximum vertical distance of 100 m (retaining ∼41%). This conservative approach accounts for the steep topography of the Powell Basin, ensuring the coupling of corresponding depth strata. PCA was run on 10,000 matched points (random subsampling with fixed seed), excluding velocity components because their circular dimension (0–360°) can inflate distances between similar directions. Depth and slope from ship bathymetry were correlated with the first two principal components. Hydrographic regimes were identified by K-means clustering to the first two principal components, with the optimal number of clusters determined via silhouette analysis. We selected k=4 (silhouette width 0.43) to retain a distinct regime that was merged at k=3 (silhouette width 0.51). The contribution of variables to cluster delineation was quantified by combining PCA loadings with Kruskal–Wallis effect sizes (η²_H) and pairwise Dunn tests. Relationships between median densities of high-density clusters with slope, depth, and bottom temperature were visualized using locally estimated scatterplot smoothing.

### Computational framework

All statistical analyses were done in R v4.5.2^77^ within [RStudio](https://posit.co/download/rstudio-desktop). Used packages were *dplyr*, *tibble*, *tidyr*, *purrr*, *forcats*, *stringr*, *scales*, and *ggplot2* from the tidyverse^78^, *ggrepel*^79^, *data.table*^80^, *vegan*^81^, *rstatix*^82^, *iNEXT*^83^*, caret*^84^, *car*^85^, *broom*^86^, *cowplot*^87^, *FactoMineR*^88^, *factoextra*^89^, *multcompView*^90^, *mvabund*^91^, *mgcv*^92^, *terra*^93^, *tidync*^94^, *sf*^95^, *dbscan*^96^, *cluster*^97^, and *FNN*^98^. Maps and 3D visualizations were created in [QGIS](https://qgis.org/) and [Global Mapper](https://www.bluemarblegeo.com/global-mapper). Figures were finalized using [Inkscape](https://inkscape.en.softonic.com/). The complete bioinformatic code and required datafiles are available under https://github.com/matthiaswietz/LandformLife and https://zenodo.org/records/18989285, respectively.

## RESULTS and DISCUSSION

Here, we integrate benthic topography and biodiversity along the Antarctic Powell Basin flank to (i) classify seafloor landforms and the geomorphological framework, (ii) identify linkages between terrain and benthic communities at local scales, and (iii) estimate benthic standing stocks across the continental slope in the oceanographic context.

### Characteristics of seafloor landforms

From ∼5000 seafloor images, captured on the Powell Basin flank between 500 and 2200 m depth, we defined seven landform classes and their geomorphology in high resolution: terraces (*n* = 35), steep slopes (*n* = 30), ridges (*n* = 14), broad slopes (*n* = 29), flat areas (*n* = 31), depressions (*n* = 28), and sand ripples (*n* = 26) (Figure 2, Supplementary Figures S2, S4). These landforms span a gradient of structural complexity. Terraces exhibit the highest heterogeneity, featuring step-like morphology (∼1 m relief) with interspersed narrow ridges and current scours (Figure 2, Supplementary Figures S5, S6) shaped by tectonics and erosion^99^. Sand ripples are less complex, featuring alternating troughs and ridges (∼20 cm relief) shaped by bottom currents (Figure 2, Supplementary Figures S5, S7). These contrasts provide a natural framework to examine how geomorphology shapes benthic communities.

**Figure 2.**
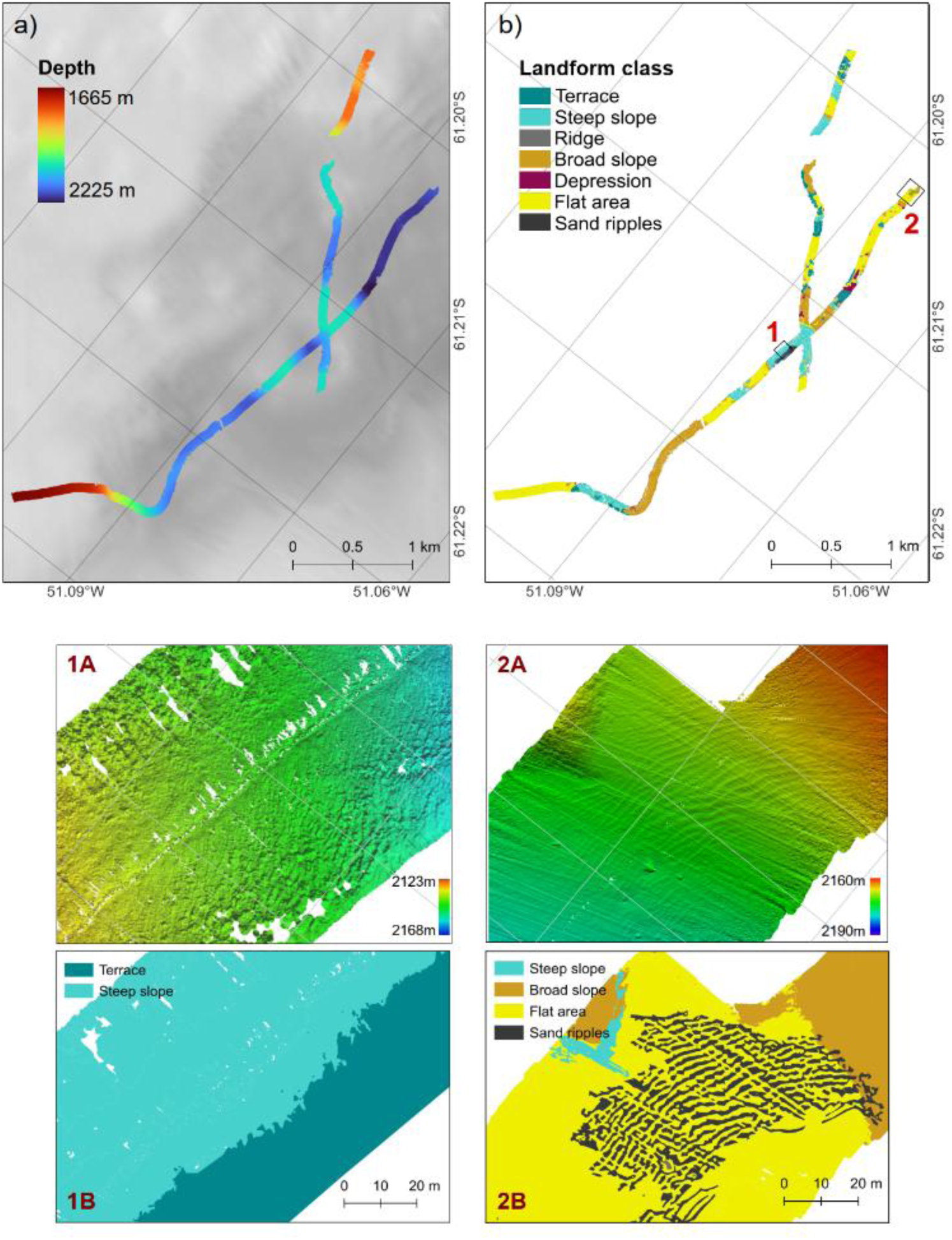
Seafloor terrain and landforms. High-resolution OFOBS bathymetry (a) and corresponding landform classification (b) along dive 69-1. Boxes labelled 1 and 2 designate areas shown in detail below, with bathymetry and landform classification for areas dominated by terraces (1A-B) and sand ripples (2A-B). Corresponding images for dives 39-1 and 81-1 are provided in Supplementary Figure S4.

Principal Component Analysis (PCA) confirmed the differentiation of landforms within their environmental setting. The first two principal components explain 37% and 24% of the total variance (Figure 3a, PERMANOVA, *p* < 0.001). PC1 primarily relates to depth, latitude and longitude, reflecting spatial distances between OFOBS dives. PC2 captured finer-scale geomorphological variability, with the strongest contributions by slope and roughness (Supplementary Table S6). Consistent with this, terraces and steep slopes – featuring significantly higher slope and roughness than other landforms – were clearly separated along this axis (Figure 3c; Dunn’s test, adjusted *p* < 0.001). Slope and roughness are highly correlated (ρ = 0.9) and jointly define surface complexity, making them the primary discriminators between landforms. To avoid collinearity, we retained only slope in subsequent analyses (see Methods). Consequently, reported slope effects capture both surface steepness and texture; features that commonly promote habitat heterogeneity and biodiversity^12–14^.

**Figure 3:**
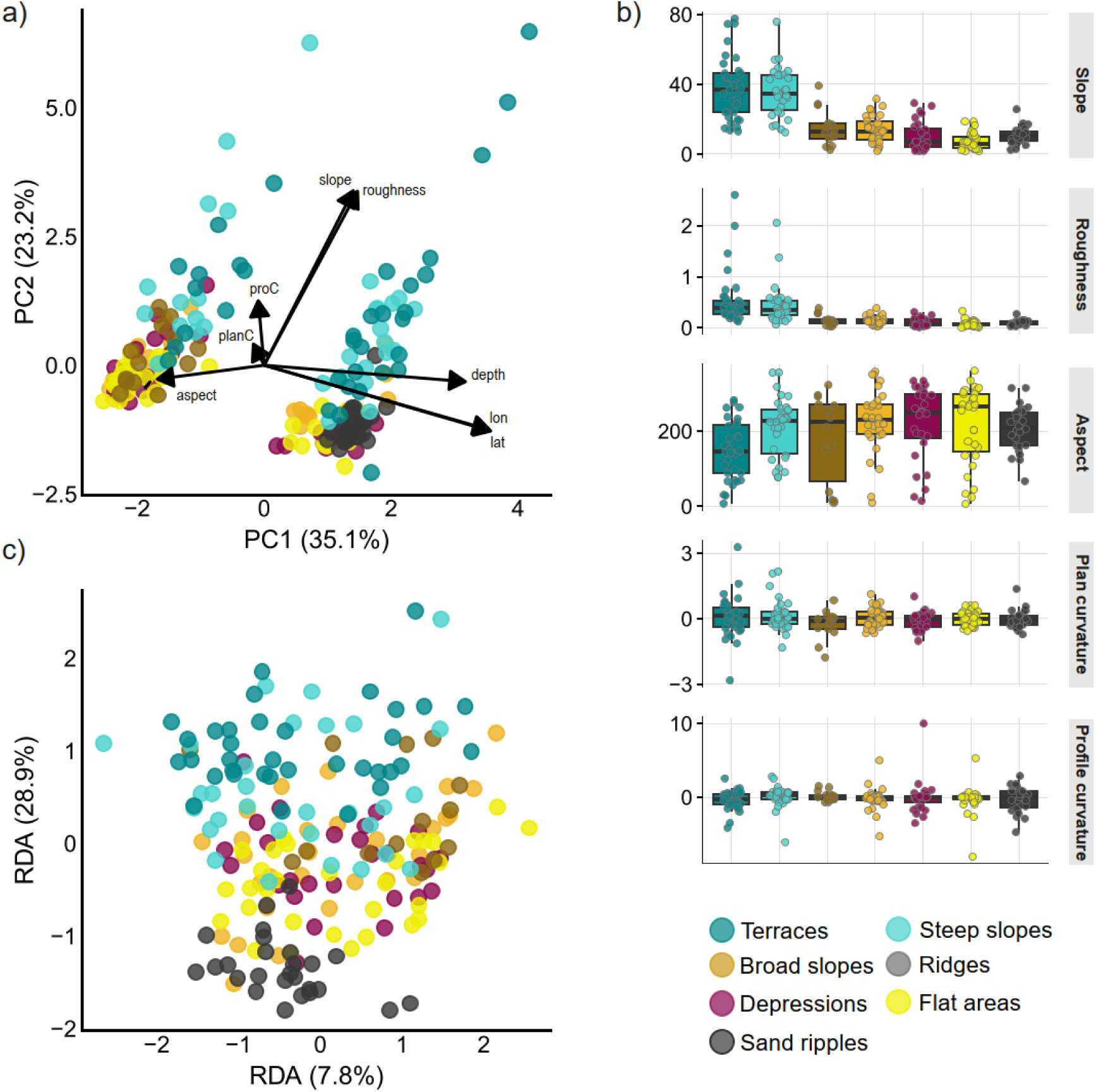
Environmental and community gradients between landforms. a) Principal Component Analysis (PCA) of terrain attributes, depth, latitude, and longitude; with samples colored by landform. b) Terrain attributes per landform. c) Redundancy analysis (RDA) of benthic community composition constrained by terrain attributes, depth, latitude, and longitude; with samples colored by landform.

### Density, diversity, and environmental drivers of benthic life

Benthic communities closely tracked this environmental framework. We quantified 10 taxa characteristic of the Antarctic benthos: corals, demosponges, glass sponges, ophiuroids, sea pens, bryozoans, anemones, asteroids, echinoids, and crinoids (Supplementary Table S5, Supplementary Table S7). Geomorphology, depth, and site jointly shape their diversity and distribution (Figures 3–4). GAMs and RDA consistently identified landforms as a major driver of benthic densities and composition (*p* < 0.001), with additional contributions from vertical and spatial gradients (GAM: *p* ≤ 0.01; RDA: *p* < 0.001). Notably, community composition was more coherently structured than densities, with RDA explaining 37% compared to 11% by GAMs, indicating a stronger geomorphological control on assemblage structure.

**Figure 4:**
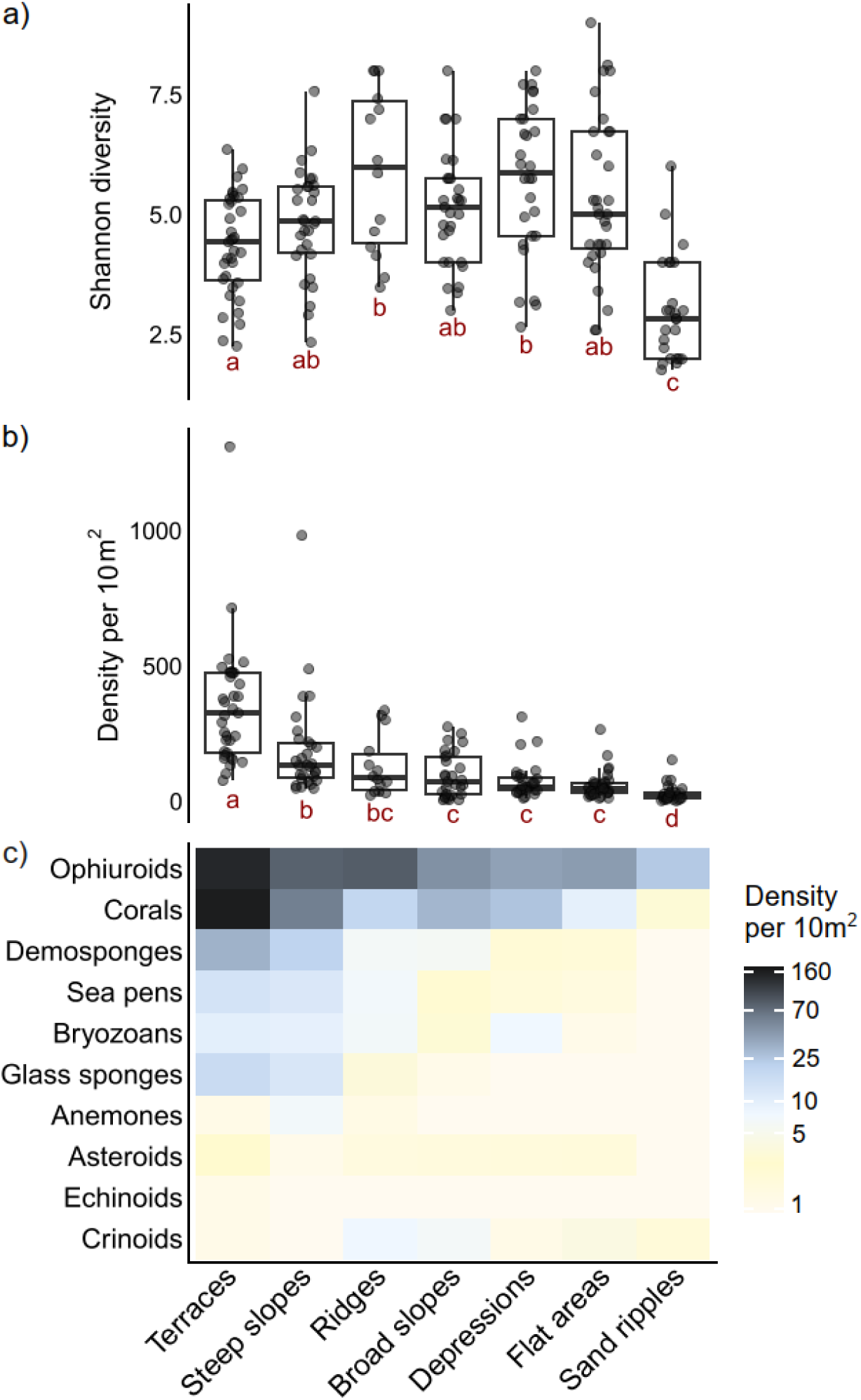
Benthic life on landforms. Alpha-diversity **(a)**, total densities **(b)**, and community composition **(c)**. Significant differences are illustrated via compact letter display: unique letters illustrate significant differences to all other landforms, while groups sharing a letter have no difference.

The role of terrain variables becomes evident when landforms are excluded. In a terrain-only GAM, slope, aspect (*p* < 0.001), and profile curvature (*p* = 0.038) are significant predictors of density (Supplementary Table S8). For RDA, variance partitioning shows that terrain attributes alone explain 1% of compositional variation, but share 10% with landforms (Supplementary Figure S8, Supplementary Table S9). Hence, terrain effects are primarily expressed through landform structure.

Ecologically, this structure is expressed in highest densities and distinct assemblages on terraces and steep slopes, with enrichment of corals, demosponges, glass sponges, ophiuroids, and sea pens (Figure 5, Supplementary Figure S9; Dunn’s test, adjusted *p* < 0.001). Terraces show characteristic biological signatures, including co-occurrence of large erect sponges and sea pens (Supplementary Figure S6). Other landforms are less differentiated, including more bryozoans on broad slopes compared to depressions, and more sea pens in depressions compared to ridges (Supplementary Figure S9). Sand ripples exhibited lowest densities and diversity, with contrasting distribution between ripple crests and troughs likely reflecting the influence of currents (Supplementary Figure S7). Ophiuroids were abundant across all landforms, highlighting their numerical dominance around the Peninsula^34^.

**Figure 5:**
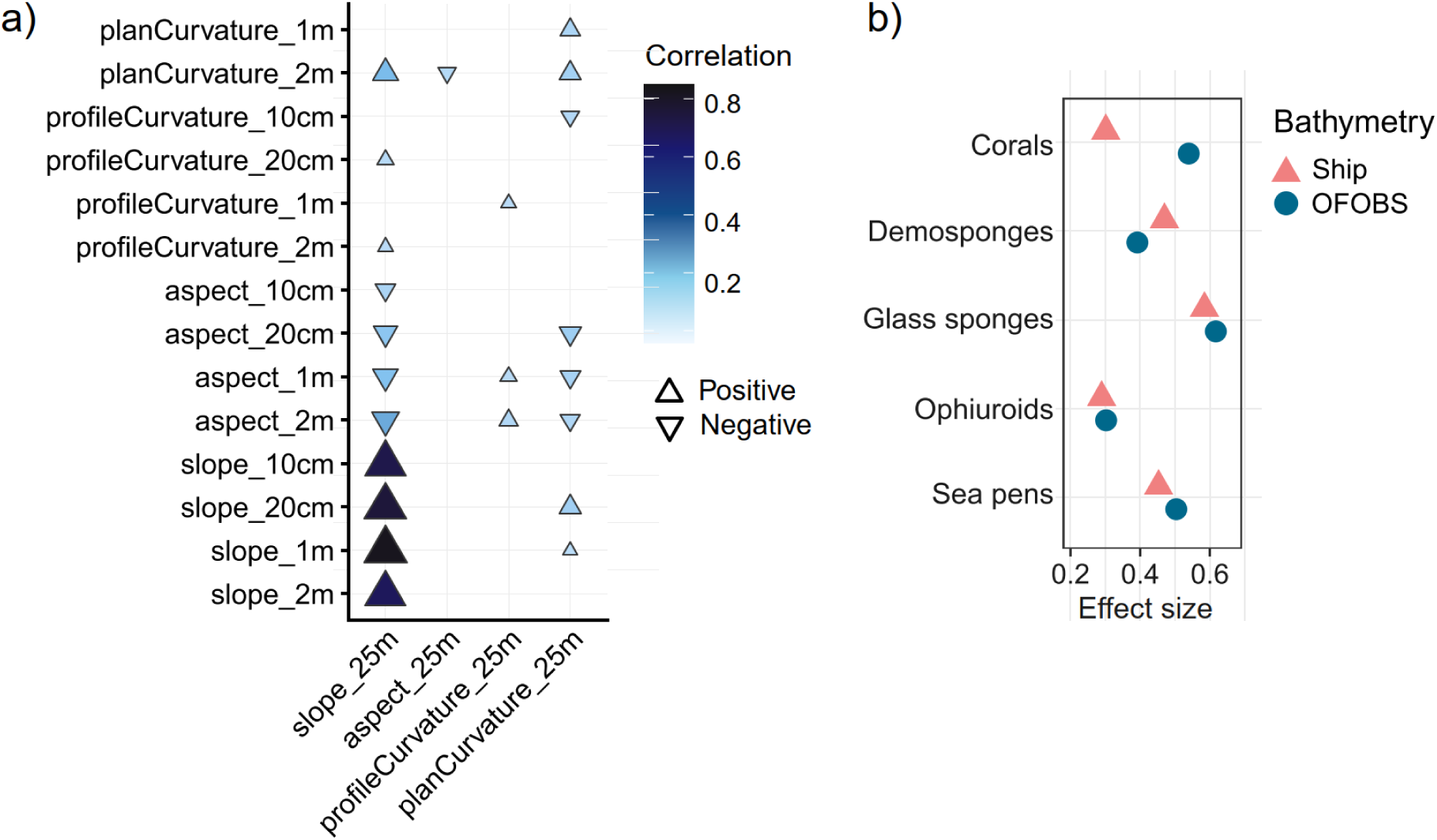
Connection between OFOBS and ship bathymetry. **a)** Pairwise Spearman’s correlations between terrain attributes across bathymetric resolutions. **b)** Effect sizes when modelling taxon densities with slope across resolutions. Only taxa and predictors with significance in both models and consistent directionality are shown.

Together, the results extend previous Weddell Sea studies^32,33,43^ by resolving benthic structuring in high geomorphological resolution. Higher densities and distinct assemblages on steep terrain highlight how topographic heterogeneity concentrates benthic biodiversity, consistent with patterns in the global deep-sea^8–11^ linked to near-bottom hydrodynamics and internal waves^15–17^. The enrichment of corals and sponges – indicator taxa for Vulnerable Marine Ecosystems (VMEs)^100^ – indicates that disturbance of these habitats has substantial ecological consequences, motivating the upscaling of local footprints to biogeographic scales.

### Biogeographic upscaling in the oceanographic context

OFOBS captures fine-scale heterogeneity, but its narrow spatial footprint limits broader biogeographic inferences. Ship bathymetry, by contrast, has extensive spatial coverage but lacks biological observations. Bridging these scales and upscaling local observations requires environmental predictors that are transferable across data sources and spatial resolutions. To address this, we compared terrain attributes derived from OFOBS and ship bathymetry, and tested whether benthic densities can be modelled using scale-independent predictors. In other words: can high- and coarse-resolution bathymetry consistently explain benthic life?

Slope was highly consistent across bathymetric resolutions (Figure 5a), whereas aspect and curvatures had weak or no correlations, indicating sensitivity to resolution and local variability. Building on this, GAMs showed that the densities of corals, demosponges, glass sponges, ophiuroids, and sea pens responded consistently to slope across resolutions (Figure 5b; Wald = 14.0 for OFOBS / 10.4 for ship, *p* = 0.001). In addition, depth was a highly significant predictor (Wald = 9.4 for OFOBS / 10.2 for ship, *p* ≤ 0.001).

Based on these robust relationships, we used these taxa and variables for regional upscaling. The resulting biogeographic map estimates a standing stock of ∼96 billion individuals across the 7,400 km² study area (Figure 6), corresponding to a mean density of ∼10 per m^2^ consistent with other Antarctic Peninsula regions^101^. The explained deviance ranged from 14–34% across taxa; as expected for spatial models of patchy benthic communities^32,34,102^. Fine-scale biological variability is hence only partially resolved at regional scales; however, estimates were stable (standard error ≈ 2,600,000).

**Figure 6.**
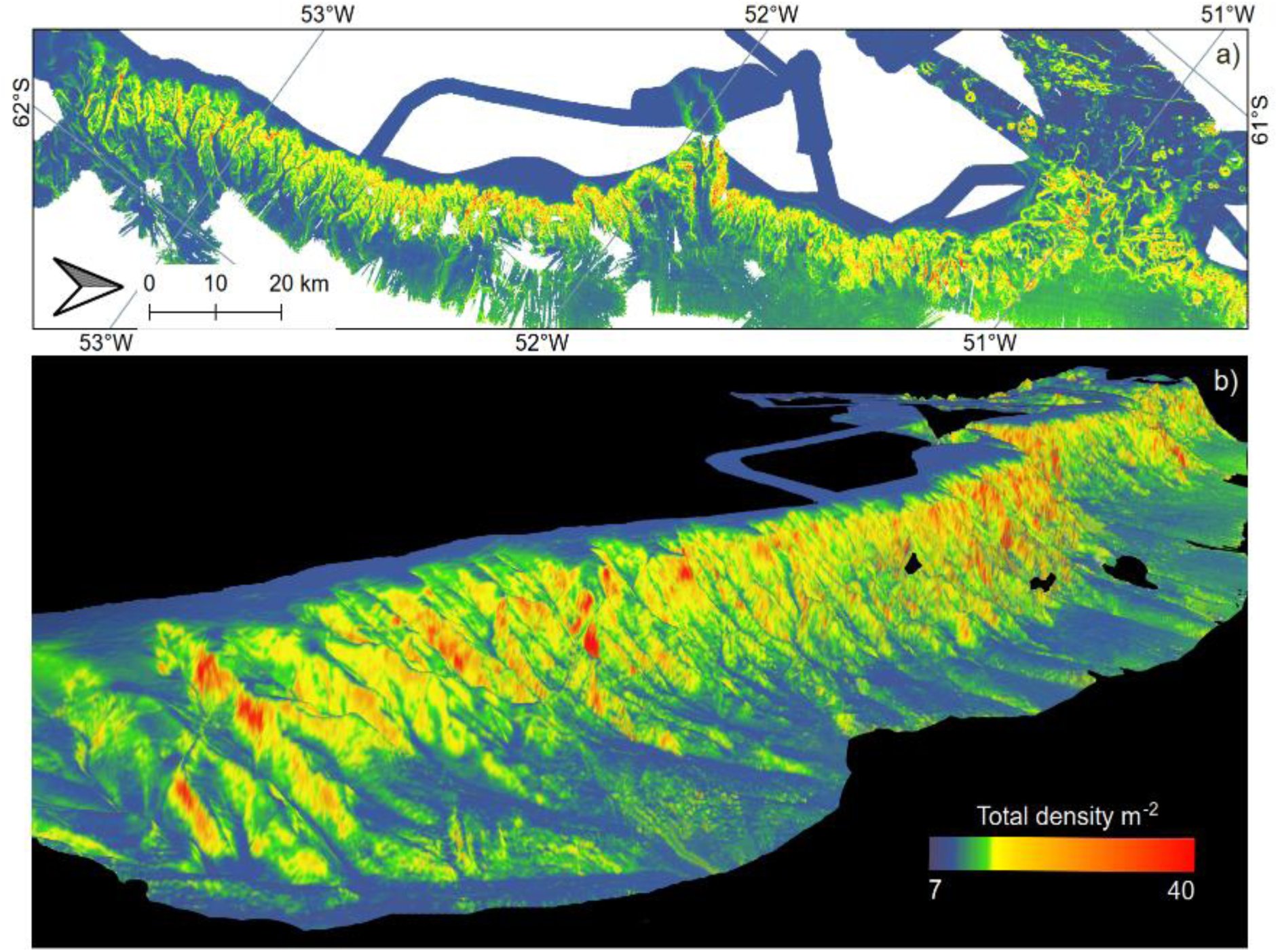
Biogeographic modelling. Predicted summed densities of corals, demosponges, glass sponges, ophiuroids, and sea pens in two-dimensional **(a)** and three-dimensional **(b)** perspectives along the Powell Basin flank.

The predictability of biodiversity through coarser bathymetric data promotes marine monitoring and management, particularly in remote regions where biological sampling is sparse or challenging^103,104^. Predictive models can translate these insights into conservation strategies^105,106^; especially in benthic systems with persistent terrain features^107^. Nonetheless, the modelling of complex biology based on restricted empirical data has limitations^50^. Slope responses may vary by site-specific influences unresolved here, such as glaciological history or shelf processes^108,109^. Furthermore, the broad taxonomic groups used for upscaling encompass numerous species with distinct niches and physiological adaptations^110,111^. Therefore, our estimates are first-order approximations. Future in-situ mapping on larger scales, leveraging autonomous surveys, automated annotation and modelling^112,113^, will help refine biogeographic conclusions.

Seafloor landforms provide the structural scaffold for benthic life, modulating local hydrodynamics within the broader oceanographic setting. To place the biogeographic upscaling into this context, we integrated 22 years of hydrographic model data^76^, matched three-dimensionally to the study area. Multivariate clustering delineated four hydrographic regimes (Figure 7a; Supplementary Table S10) with significant differences in bottom temperature, temperature in the overlying water, stratification, sea-ice concentration, and chlorophyll-a (Dunn’s test, adjusted *p* < 0.001). Projecting benthic data onto these regimes revealed significant differences in depth, slope, and predicted densities (effect sizes η²H = 0.58, 0.29, and 0.1, respectively). Decadal water mass structure thus covaries with seafloor topography and biogeography.

**Figure 7.**
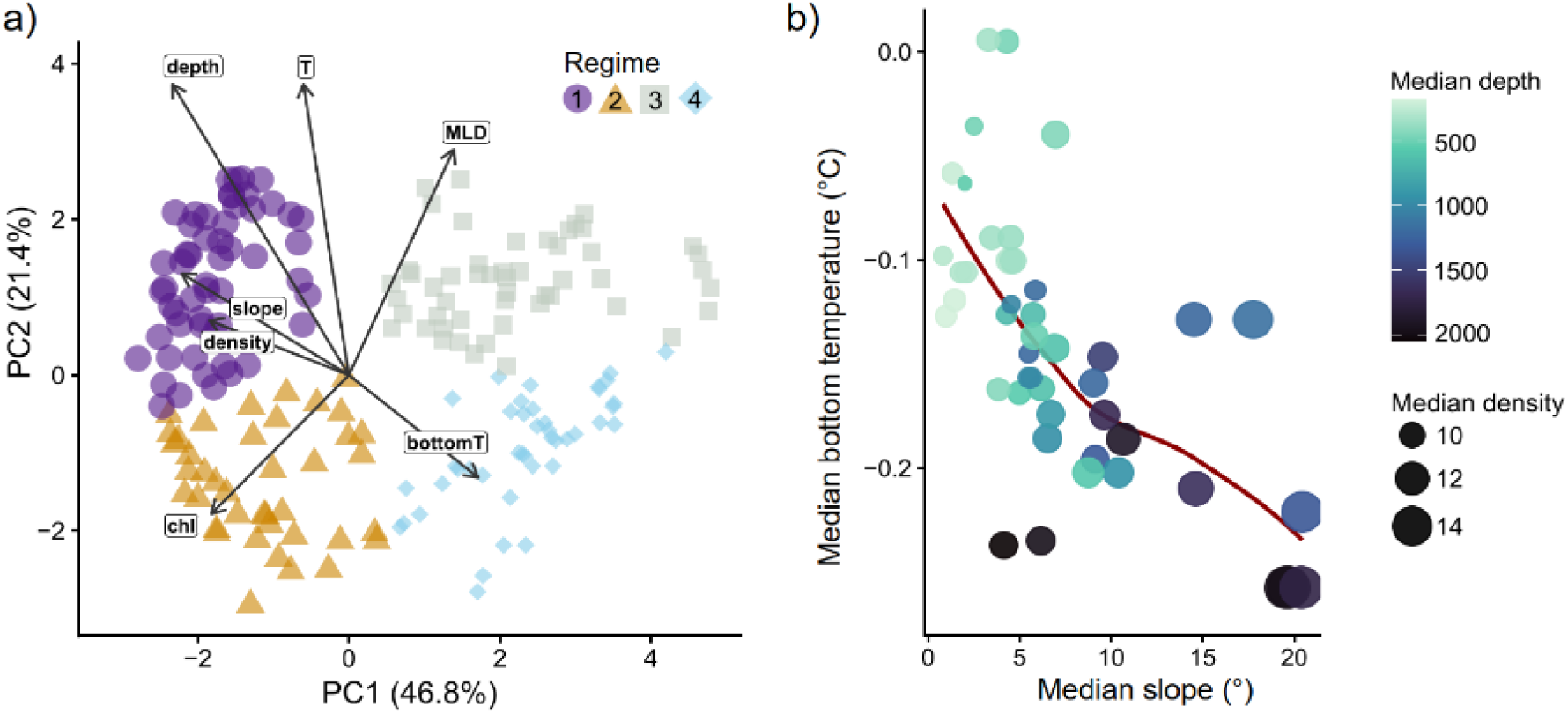
Bathymetric, hydrographic, and biological connections. **a)** Principal component analysis and K-means clustering of decadal oceanographic model data delineates four hydrographic regimes. The arrows for slope and depth correspond to their correlation with the two first principal components. Dot size indicates median predicted densities of corals, sponges, ophiuroids, and sea pens projected onto these regimes, ranging from 12.0 (regime 1) to 9.3 (regime 4). **b)** Increasing densities within high-density clusters along the gradient of bottom water temperature, slope, and depth, illustrated by the red loess line. T: potential temperature in the water column, bottomT: potential temperature near the seafloor, Ice: sea-ice concentration, MLD: mixed-layer depth, chl: chlorophyll.

### Biodiversity hotspots

Deep-sea benthos often exhibits local patchiness, yet the distribution and environmental controls of biodiversity maxima remain elusive^114,115^. We identified biological peaks by aggregating grid cells within the top 10% of predicted densities – resulting in 47 high-density clusters and ultimately four spatially coherent hotspots. Across hierarchical levels, densities peak within a consistent environmental envelope: steep terrain with >10° slope, depths >1,400 m, and bottom temperatures below −0.15 °C (Figure 7b). Consistently, “hot” grid cells are enriched in the cold and deep hydrographic regime (chi-squared test, *p* < 0.001).

Hotspots constitute 0.5% of the study area (∼33 km²) with median depths of 1,770 m and median slopes of 35° (Figure 8a), including deeply incised canyons (Figure 8b). Hotspots host 2.5-fold higher benthic densities, with enrichment factors between 1.3 (sea pens) and 3.0 (corals and glass sponges). Density maxima align with the pathway of Weddell Sea Deep Water (WSDW) at its interface with Weddell Sea Slope Water (WSSW), yielding distinct bottom-water conditions^60^. Peaks around steep canyons, through which WSDW is likely exported into the Bransfield Strait^116,117^, indicate channeling, upwelling, and mixing processes^118–120^. Such hydrodynamics are known to promote benthic life through elevated carbon and oxygen levels^121–123^ and facilitated dispersal of larvae^124^.

**Figure 8:**
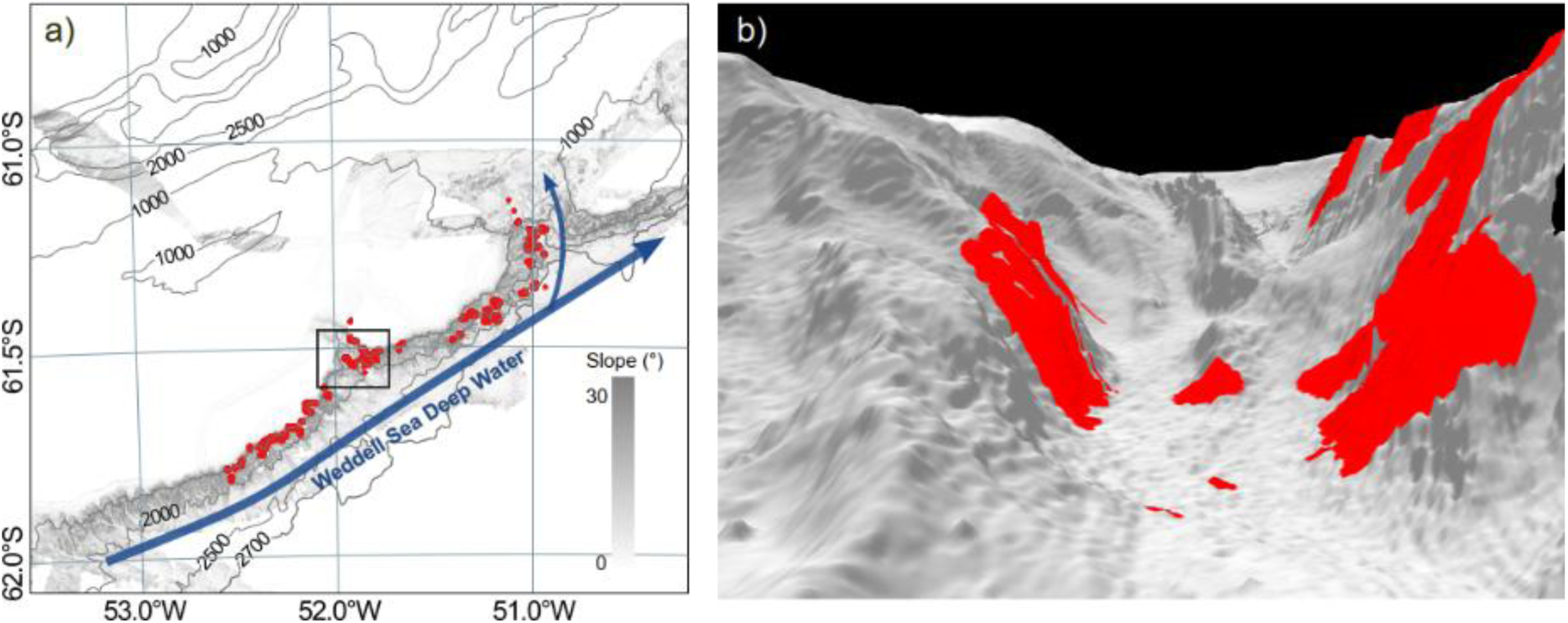
Benthic hotspots and their water mass alignment. **A:** Predicted benthic hotspots (red) along the flow of Weddell Sea Deep Water (blue). **B:** Focused view of a “hotspot canyon” indicated by the rectangle in a). The terrain bathymetry is depicted using hillshade to enhance topographic visualization.

Notably, temperatures near the seafloor and in the overlying water are decoupled around hotspots (Figure 9b): the mean difference increases from non-hotspots (0.08°C) to high-density clusters (0.14°C) to hotspots (0.21°C). This divergence remained significant when accounting for depth and slope (GAM*; p* < 0.001), ruling out simple terrain effects. Spatial mismatches are also unlikely, as temperature data were stringently mapped to the cruise coordinates (≤10 km horizontal and ≤100 m vertical distance; see Methods). Instead, hotspots likely coincide with localized hydrographic transitions along steep walls, where cold WSDW near the seafloor is overlain by relatively warmer waters^125^. Similar processes enhance cross-slope hydrography in other Antarctic canyons^126^. At the same time, bottom temperature models might be overestimated relative to in-situ observations^127^. Hence, the temperature decoupling probably reflects a combination of physical processes and limited model representation over steep, incised terrain. Density maxima also coincided with shallower mixed layers and higher sea-ice cover (Figure 9a; Dunn’s test, adjusted *p* < 0.05), indicating distinct cryo–pelagic–benthic coupling. Within this element of the polar biological carbon pump, downward organic matter pulses provide an energetic link between surface and benthos^128–130^. However, surface signatures may also reflect coincidental circulation and sea-ice dynamics^127^.

**Figure 9:**
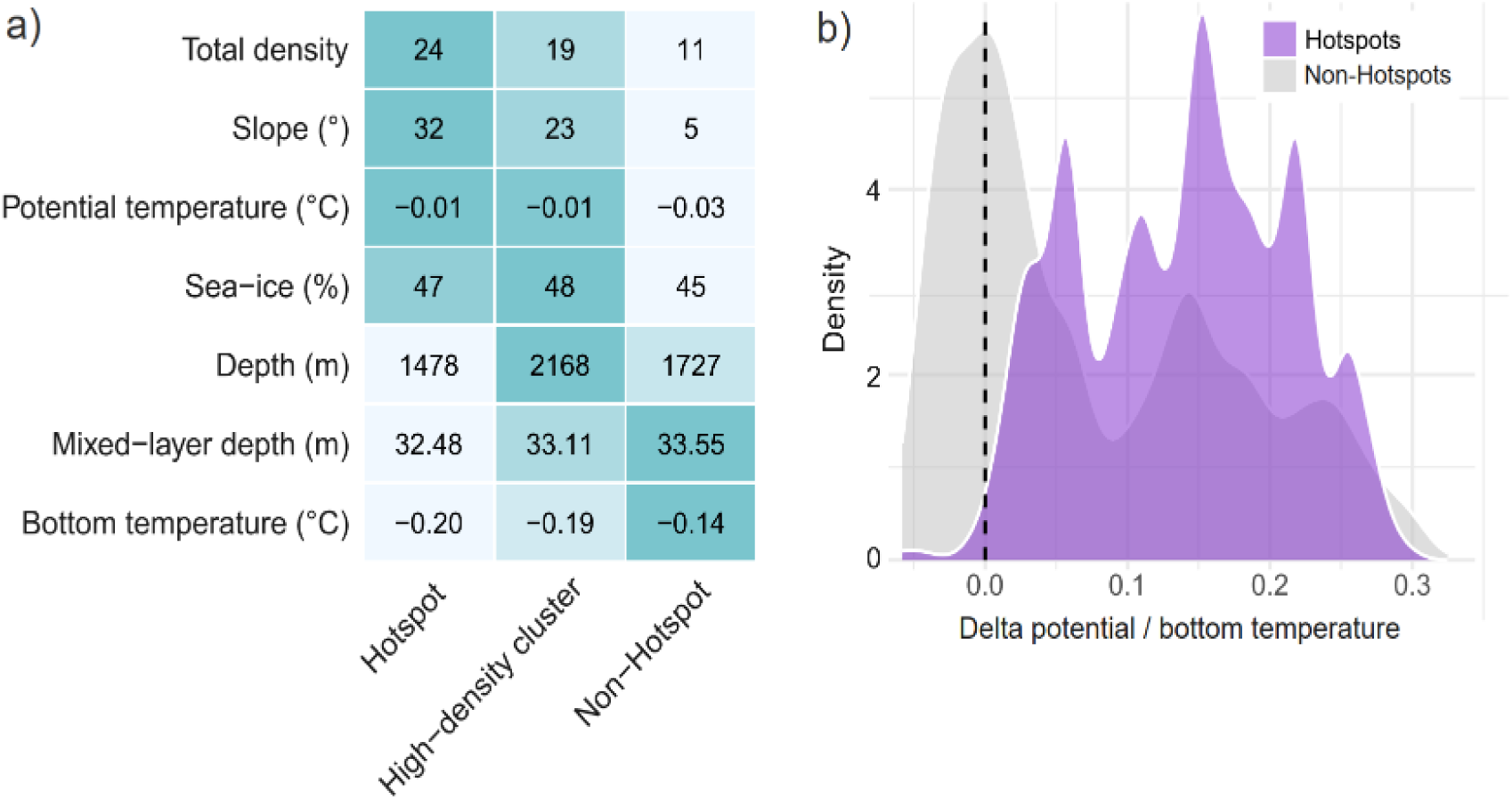
Hotspot characteristics. a) Environmental gradients between hotspots, high-density clusters, and non-hotspots, showing variables with significant differences relative to non-hotspots (Dunn’s test, adjusted *p* < 0.05) independent of spatial extent. b) Significant decoupling (increased delta) between potential temperature and bottom temperature around hotspots and high-density clusters.

### SYNOPSIS

Seafloor geomorphology and oceanography jointly shape benthic biodiversity along the Powell Basin flank. Steep terrain concentrates benthic life, while water-mass circulation and thermal regimes guide where biological hotspots occur. Benthic ecosystems are hence likely sensitive to shifting environmental regimes, given the narrow thermal tolerances of many Antarctic taxa^131–133^. Although carbon cycling was not quantified here, benthic communities are recognized carbon sinks, and changes in their distribution may influence long-term carbon cycling^134–136^. At the same time, physically stable features like steep slopes may provide environmental stability, potentially acting as refugia − a concept recently proposed for other benthic habitats and taxa^137^. The identification of key environmental controls on Antarctic benthic biodiversity establishes a framework for understanding ecosystem structure, monitoring future changes, and guiding conservation in the warming Weddell Sea^138–140^.

## SUPPLEMENTARY FIGURES

**Supplementary Figure S1.**
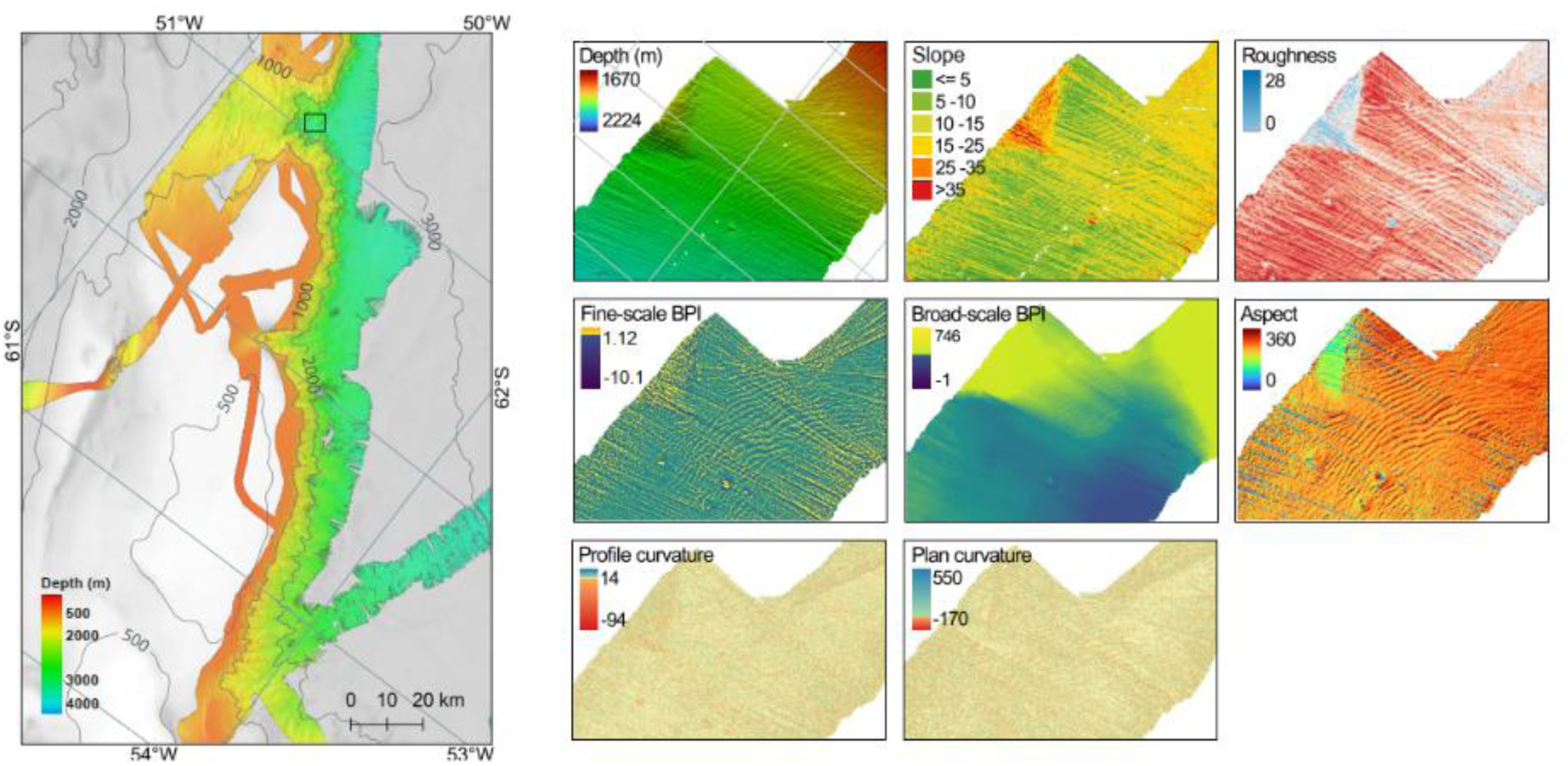
Ship bathymetry of the study area (25 m resolution) and OFOBS-derived terrain attributes (20 cm resolution) for an example section (indicated by the black rectangle).

**Supplementary Figure S2.**
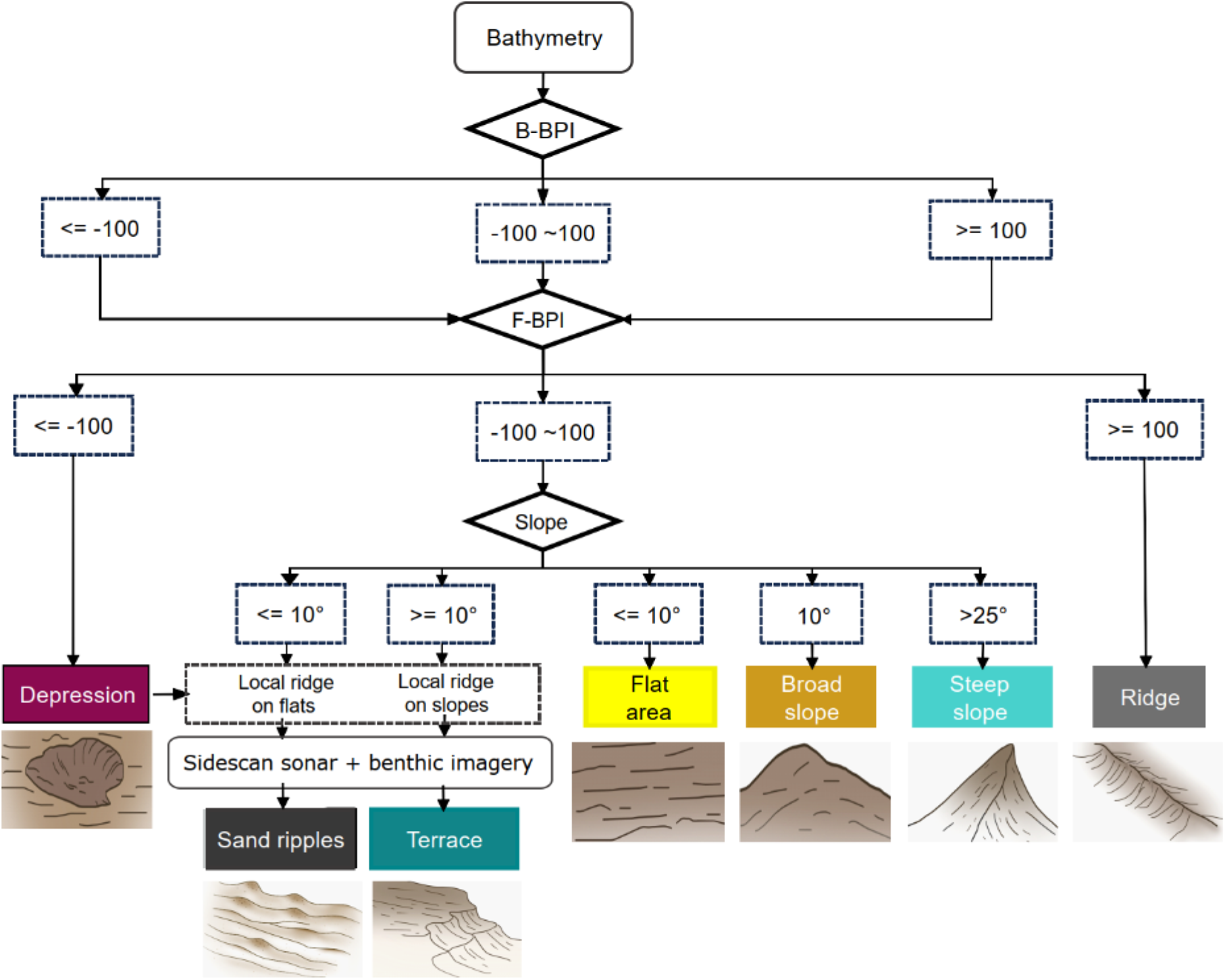
Landform classification based on fine-scale (F-BPI) and broad-scale (B-BPI) bathymetric positioning index, together with slope. Compound landforms (terraces and sand ripples) were additionally defined through characteristic spatial patterns in high-resolution bathymetry and backscatter mosaics, as confirmed using seafloor imagery. Sketches of each landform are depicted below.

**Supplementary Figure S3.**
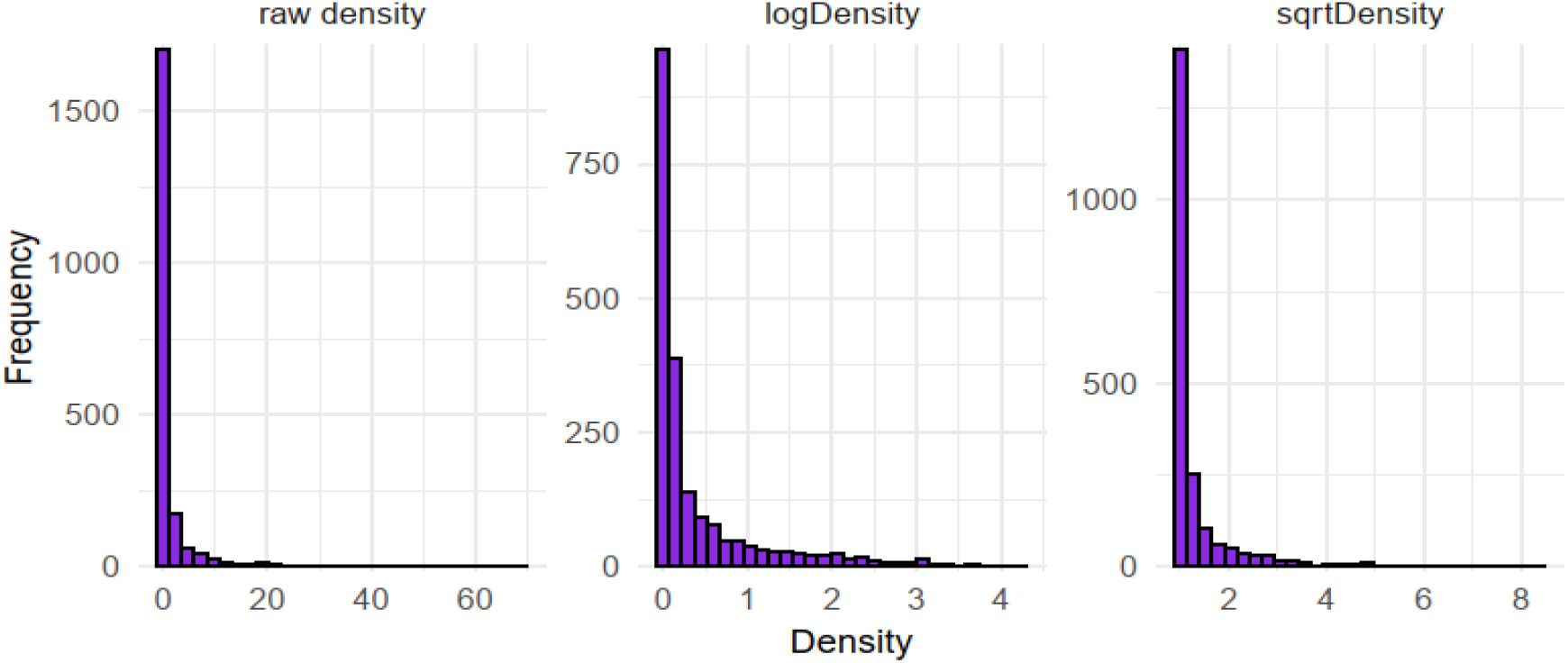
Histogram of raw, log- and square-root transformed taxon counts identified log-transform as best, and was subsequently used for assessing benthic density.

**Supplementary Figure S4.**
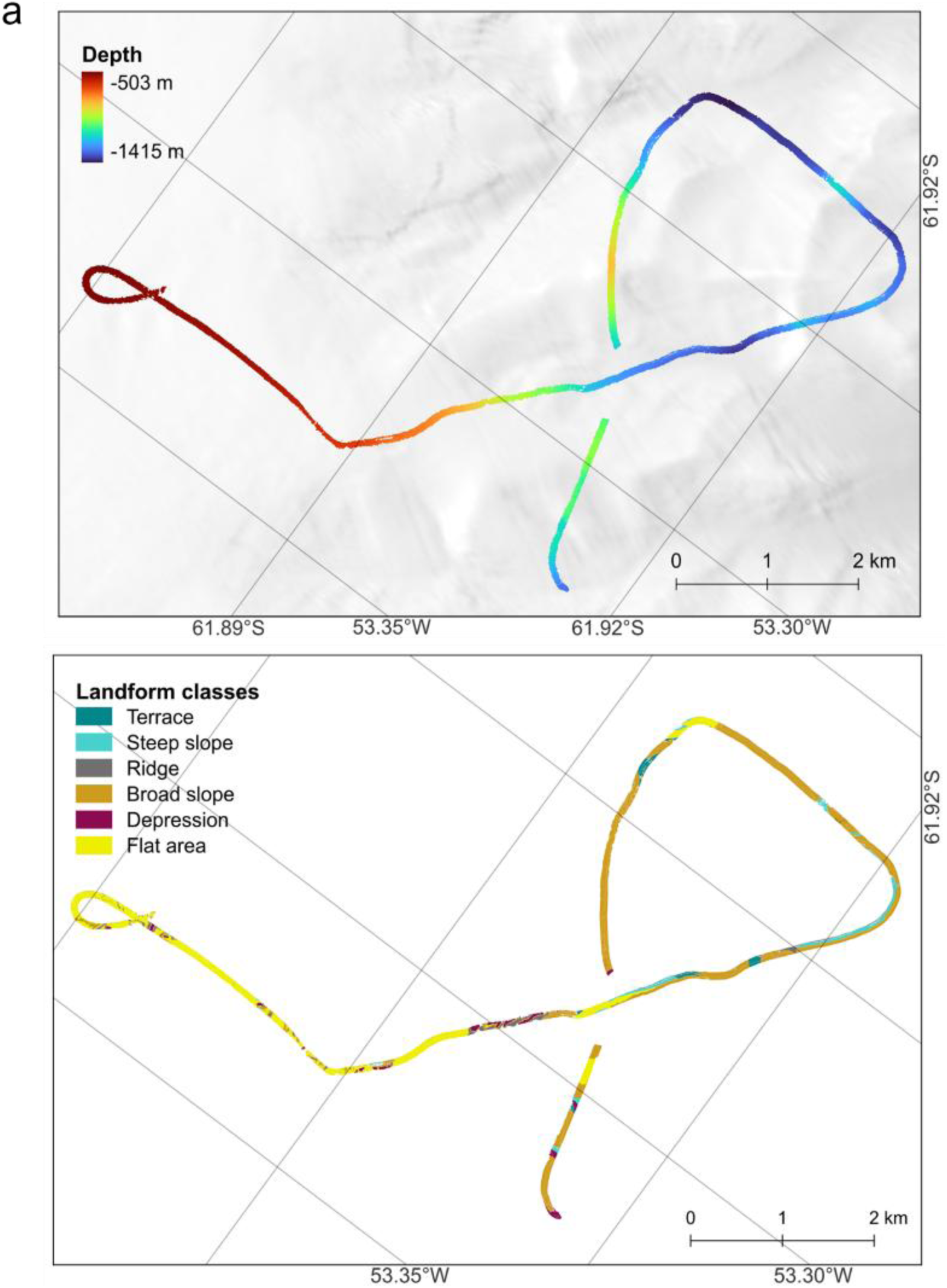
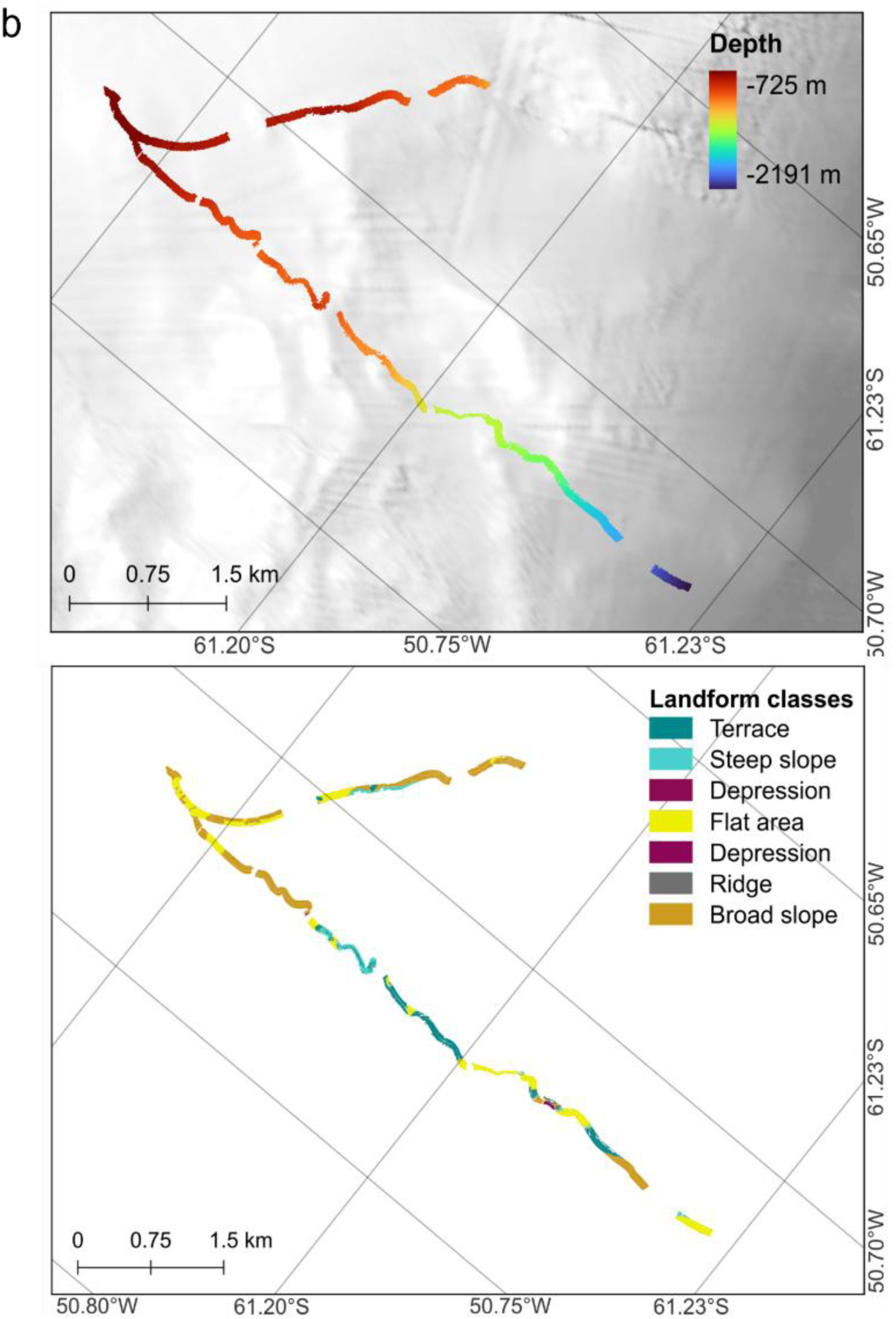
Bathymetry and landform classes for dives 39-1 (a) and 81-1 (b).

**Supplementary Figure S5:**
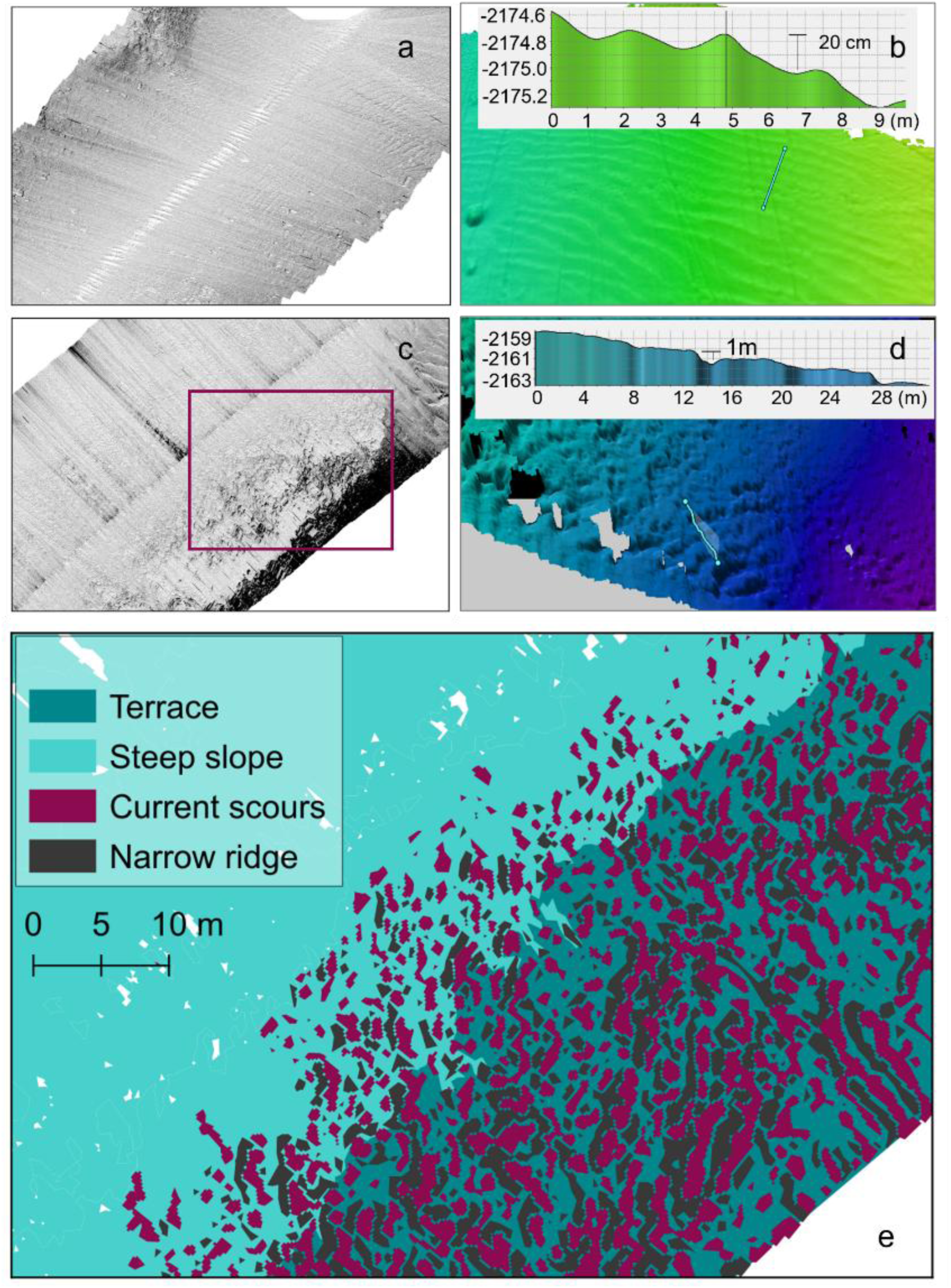
Landform terrain features. a–b) Sand ripples illustrated by 3D bathymetry and sidescan. The insert in b) depicts ripple heights along a 9 m transect. c–d) Terraces illustrated by 3D bathymetry and sidescan. The insert in d) depicts terrace step heights along a 28 m transect. e) Zoomed-in view of the red box in (c), illustrating the topographic complexity on terraces with narrow ridges and current scours.

**Supplementary Figure S6:**
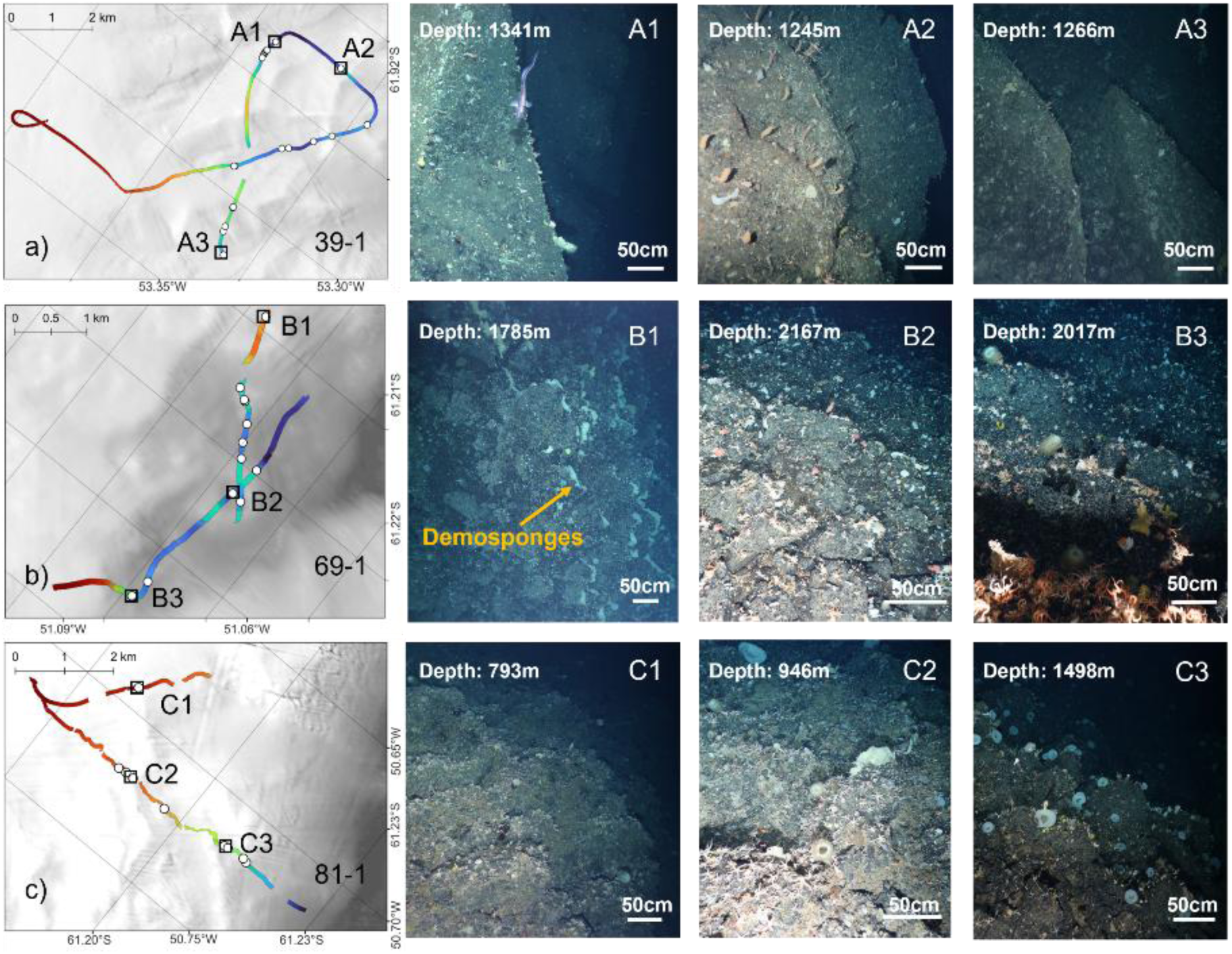
Terraces and epibenthos. Locations of terraces on OFOBS dives 39-1, 69-1 and 81-1 (white dots in panels a-c) and example images of associated benthic communities (A1-C3). Arrows indicate demosponges on terrace edges.

**Supplementary Figure S7:**
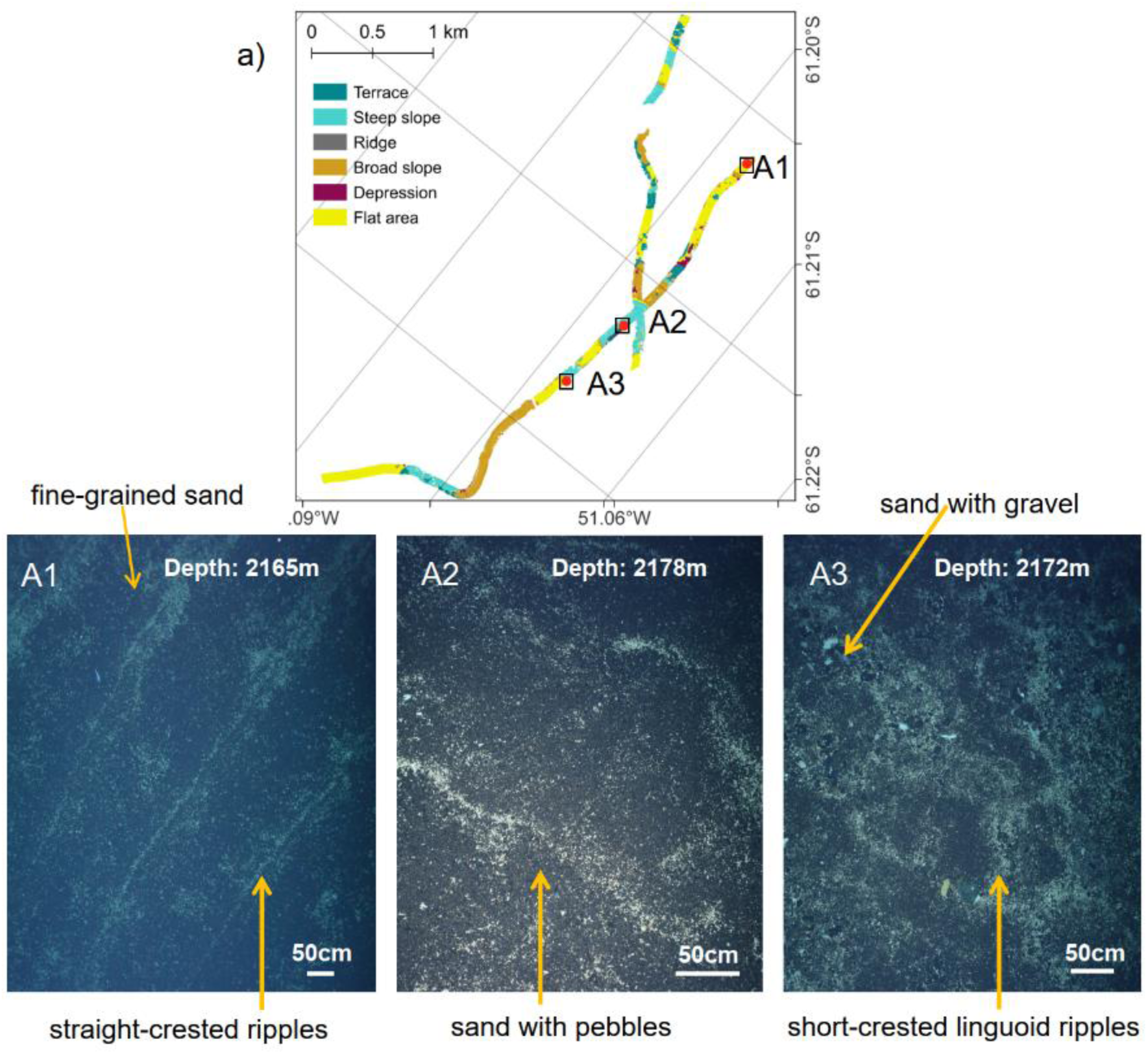
Sand ripples and epibenthos. Locations of sand ripples in dive 69-1 (a), and example images of associated benthic communities (A1-A3). A1: fine-grained sand substrate with a straight-crested ripple pattern. A2: sandy substrate mixed with pebbles. A3: sandy substrate with gravel. Ripples in fine-grained sand indicate steady, slow bottom currents whereas linguoid ripples associated with gravel and pebble patches suggest increased flow velocities.

**Supplementary Figure S8:**
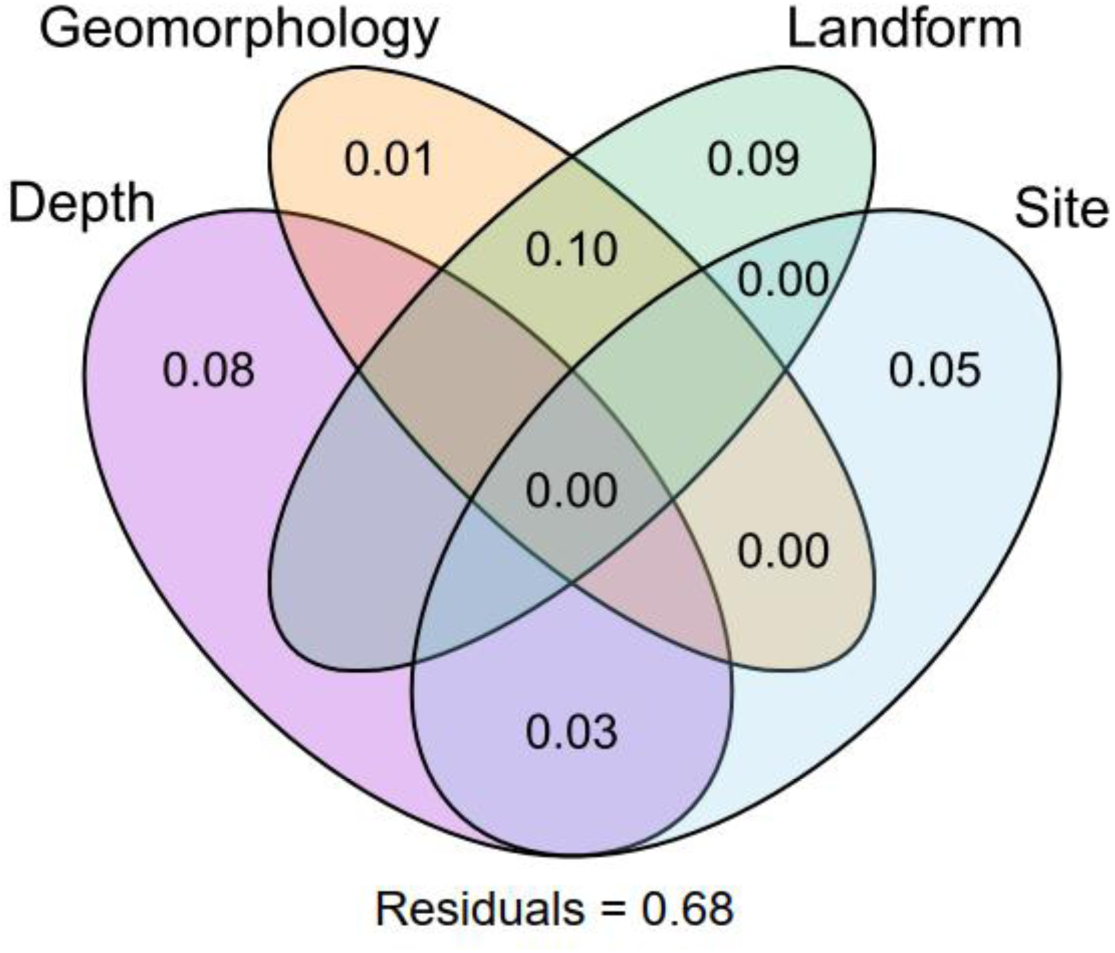
Variance partitioning of RDA predictors. Venn diagram showing the relative contribution of variable groups on community discrimination between landforms in RDA.

**Supplementary Figure S9:**
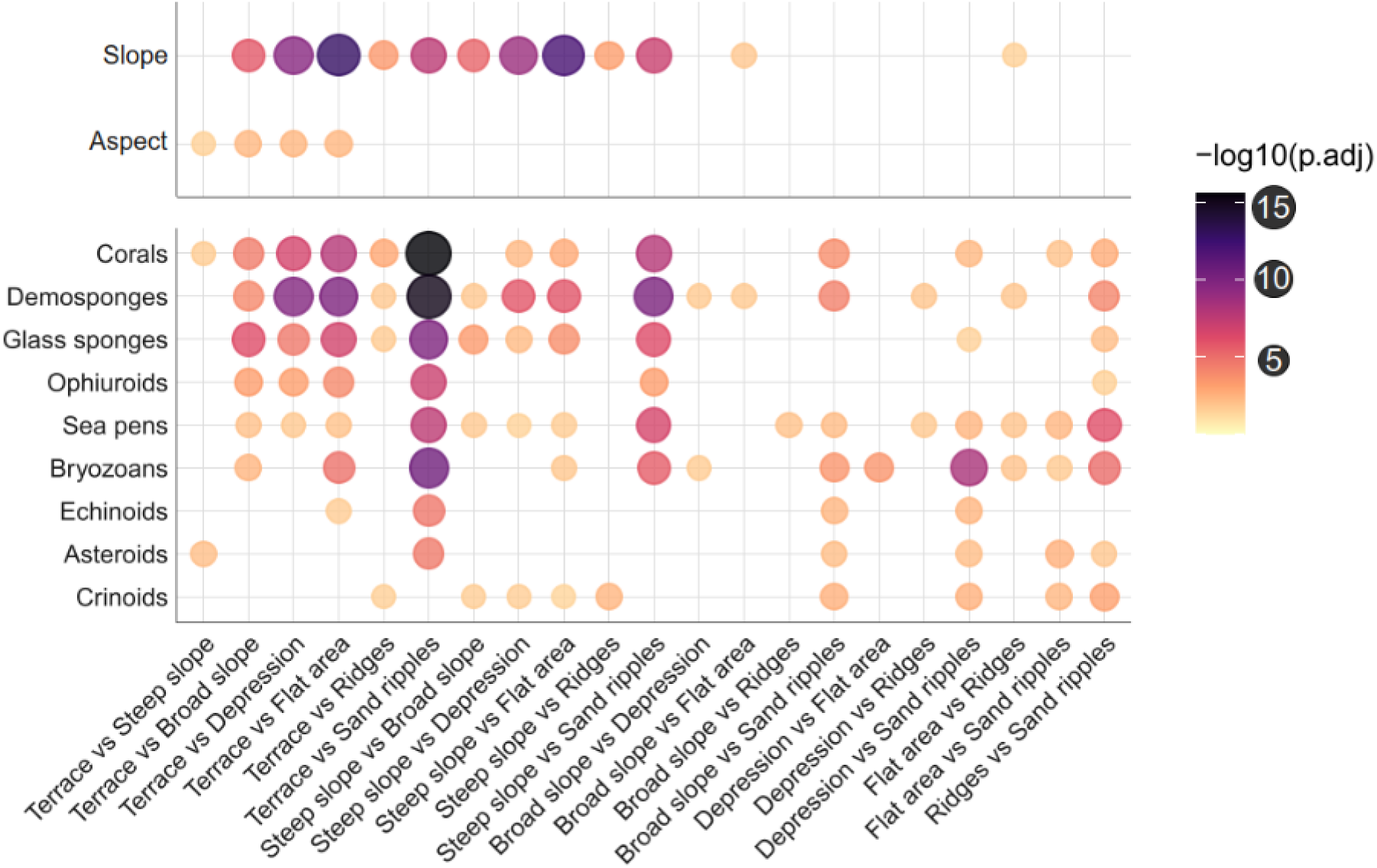
Significantly differing terrain variables (top) and taxon abundances (bottom; Terraces and Steep slopes highlighted in green shades) between landforms.

## ACKNOWLEDGEMENTS

We thank the captain, crew, and scientific team of expedition PS118 for their outstanding support, especially Laura Hehemann, Simon Dreutter, and Axel Nordhausen for operating OFOBS. Special thanks to Jan Jansen, Lili Böhringer, Santiago Pineda Metz, and Julian Gutt for their expert taxonomic advice. We thank Markus Janout for oceanographic advice and Simon Dreutter for supporting bathymetric data acquisition and processing. This work was supported by the National Natural Science Foundation of China (Grant No. 42206200) and the China Scholarship Council. MW was funded through the DFG Priority Program SPP 1158 “Antarctic Research with comparative investigations in Arctic ice areas” (Grant No. 522416631). AWI Grant AWI_PS118_09 supported AP and the OFOBS team.

## DATA AVAILABILITY

OFOBS photographs for dives 39-1, 69-1, and 81-1 are available on PANGAEA. [OFOBS](https://doi.org/10.1594/PANGAEA.930222) and [ship](https://doi.pangaea.de/10.1594/PANGAEA.916118) bathymetry are available on PANGAEA. Oceanographic and chlorophyll data were downloaded from the Copernicus Marine Environment Monitoring Service (CMEMS) [Global Ocean Physics](https://data.marine.copernicus.eu/product/GLOBAL_MULTIYEAR_PHY_001_030/description) and [Ocean Colour](https://doi.pangaea.de/10.1594/PANGAEA.916118) model products, respectively. The complete bioinformatic code and required datafiles are available under https://github.com/matthiaswietz/LandformLife and https://zenodo.org/records/18989285, respectively.

## AUTHOR CONTRIBUTIONS

MF, MW and BD conceived the study. MF and MW analyzed the data, interpreted and visualized results, and wrote the paper. AP collected the data and contributed to image evaluation. TI, BW, and NC contributed to interpretation and writing.

