## Supplemental figures for "Benthic biodiversity hotspots in the Weddell Sea: Bridging geomorphology, biogeography, and oceanography across scales"

### SUPPLEMENTARY FIGURES

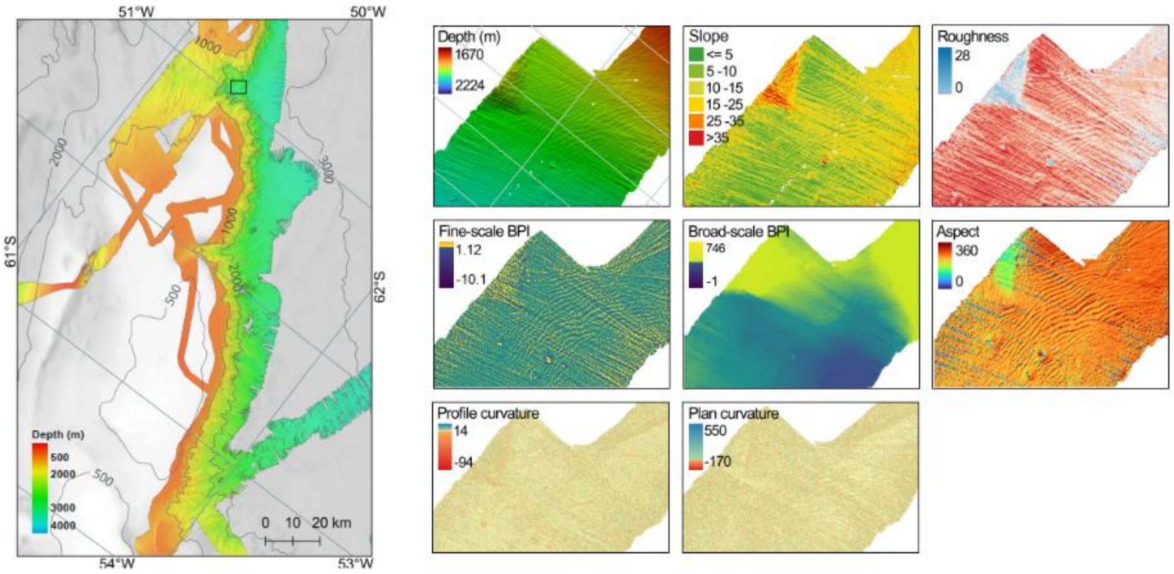


**Supplementary Figure S1.** Ship bathymetry of the study area (25 m resolution) and OFOBS-derived terrain attributes (20 cm resolution) for an example section (indicated by the black rectangle).

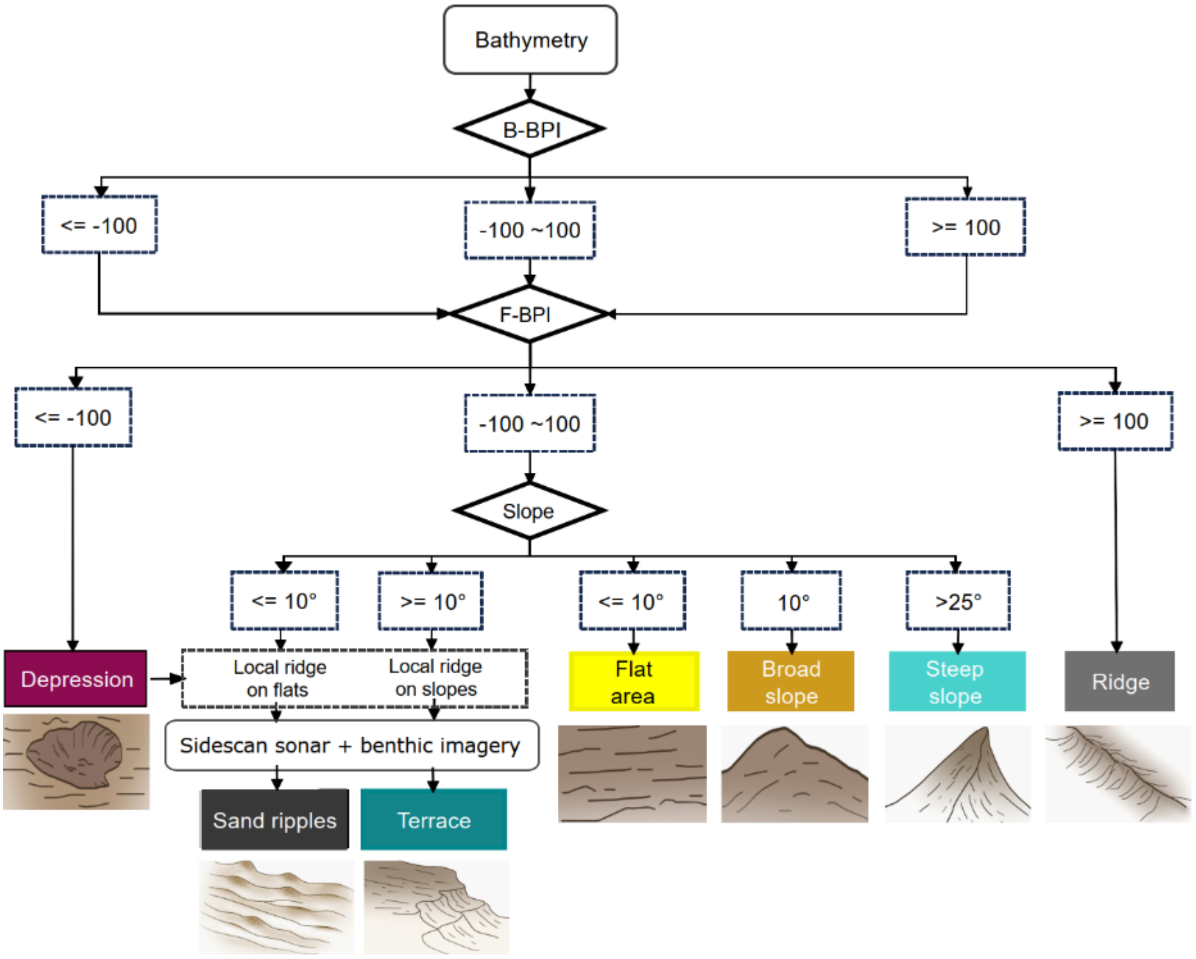


**Supplementary Figure S2** Landform classification based on fine-scale (F-BPI) and broad-scale (B-BPI) bathymetric positioning index, together with slope. Compound landforms (terraces and sand ripples) were additionally defined through characteristic spatial patterns in high-resolution bathymetry and backscatter mosaics, as confirmed using seafloor imagery. Sketches of each landform are depicted below.

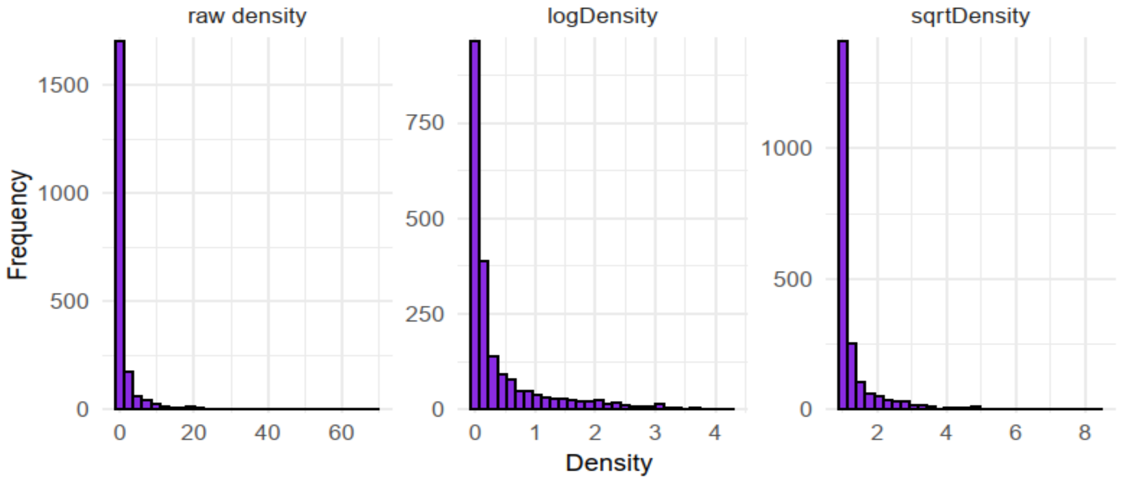


**Supplementary Figure S3.** Histogram of raw, log- and square-root transformed taxon counts identified log-transform as best, and was subsequently used for assessing benthic density.

a

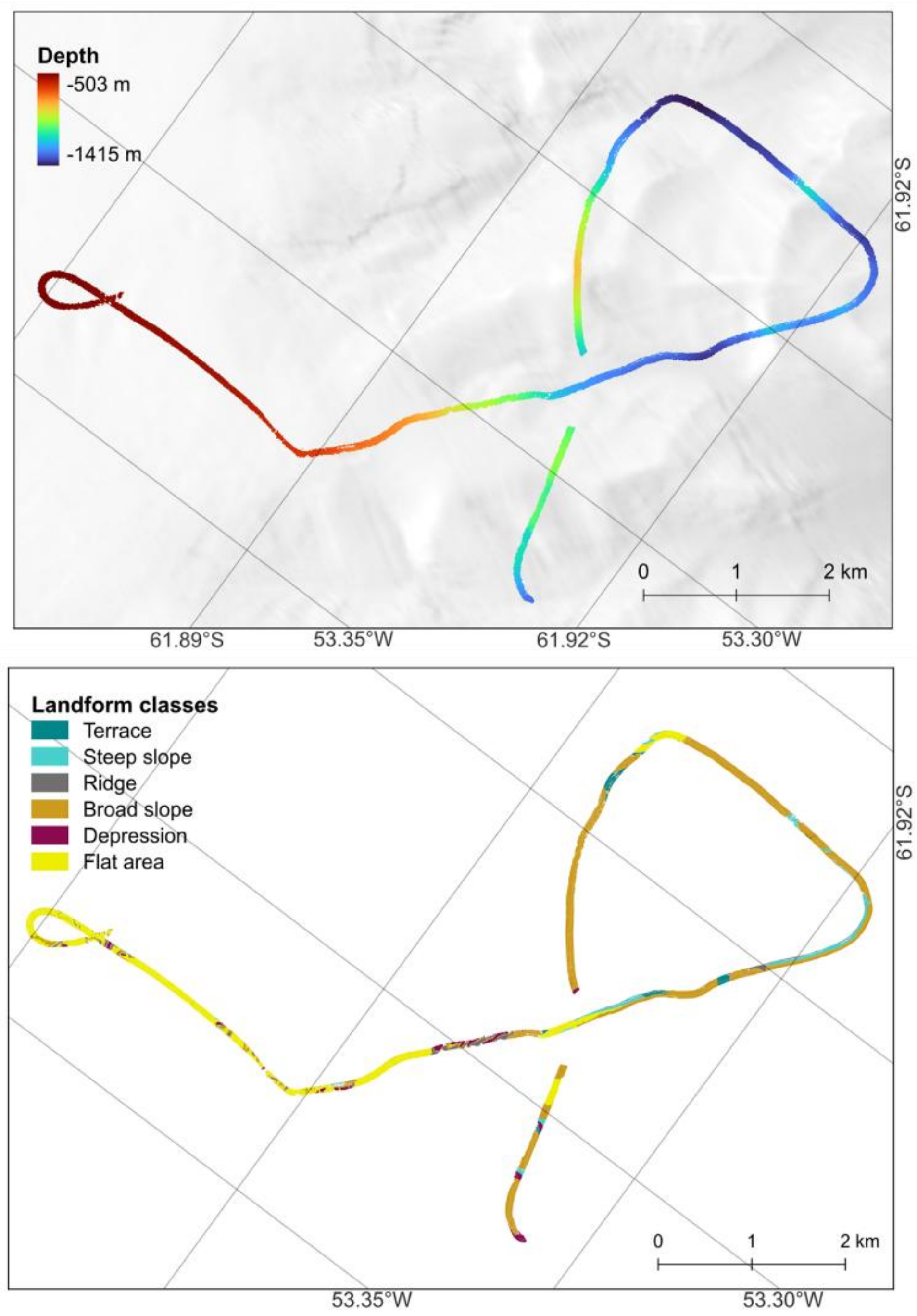

Depth
-503 m
-1415 m
61.92°S
0
1
2 km
61.89°S
53.35°W
61.92°S
53.30°W
Landform classes
Terrace
Steep slope
Ridge
Broad slope
Depression
Flat area
61.92°S
0
1
2 km
53.35°W
53.30°W

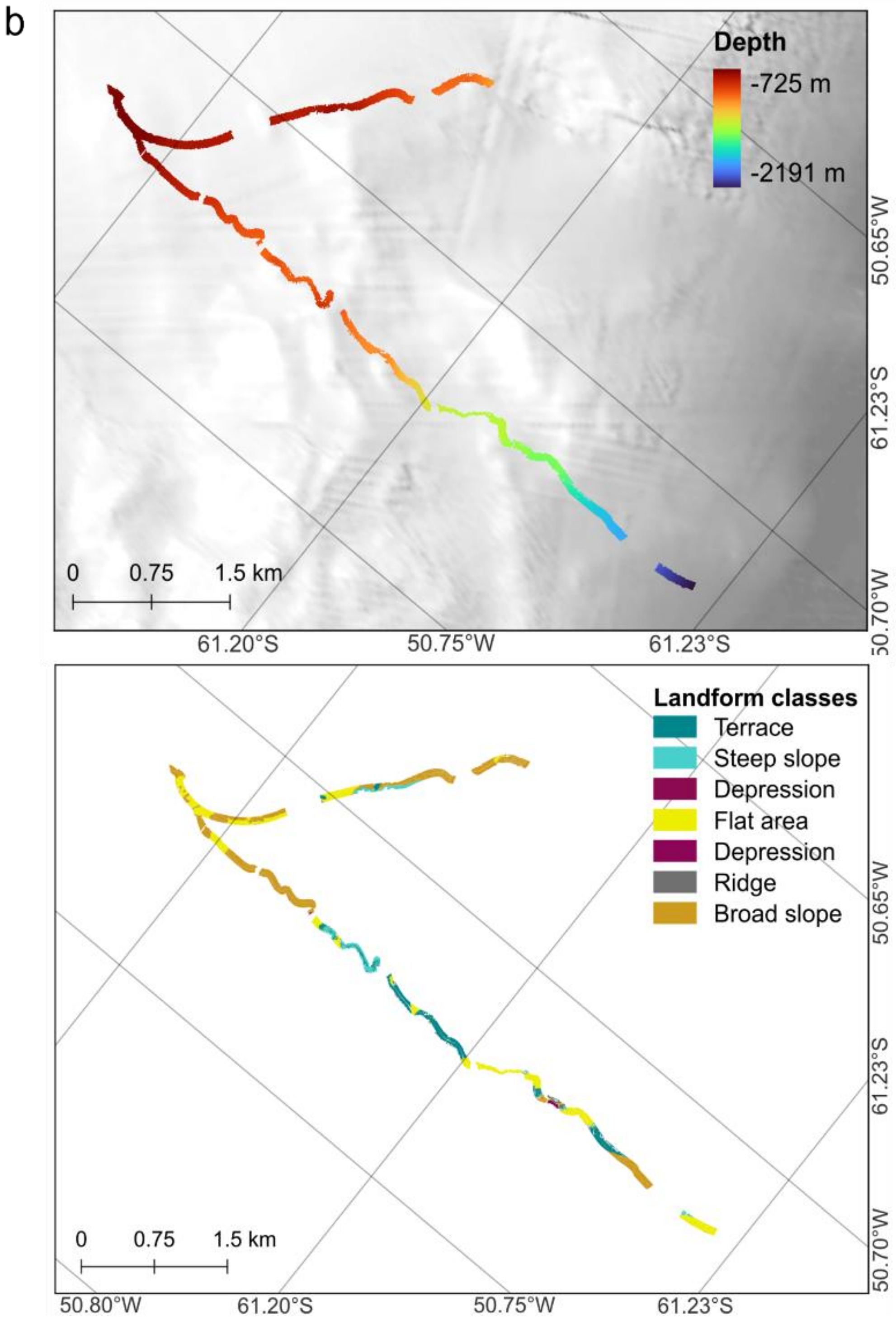


**Supplementary Figure S4** Bathymetry and landform classes for dives 39-1 (a) and 81-1 (b).

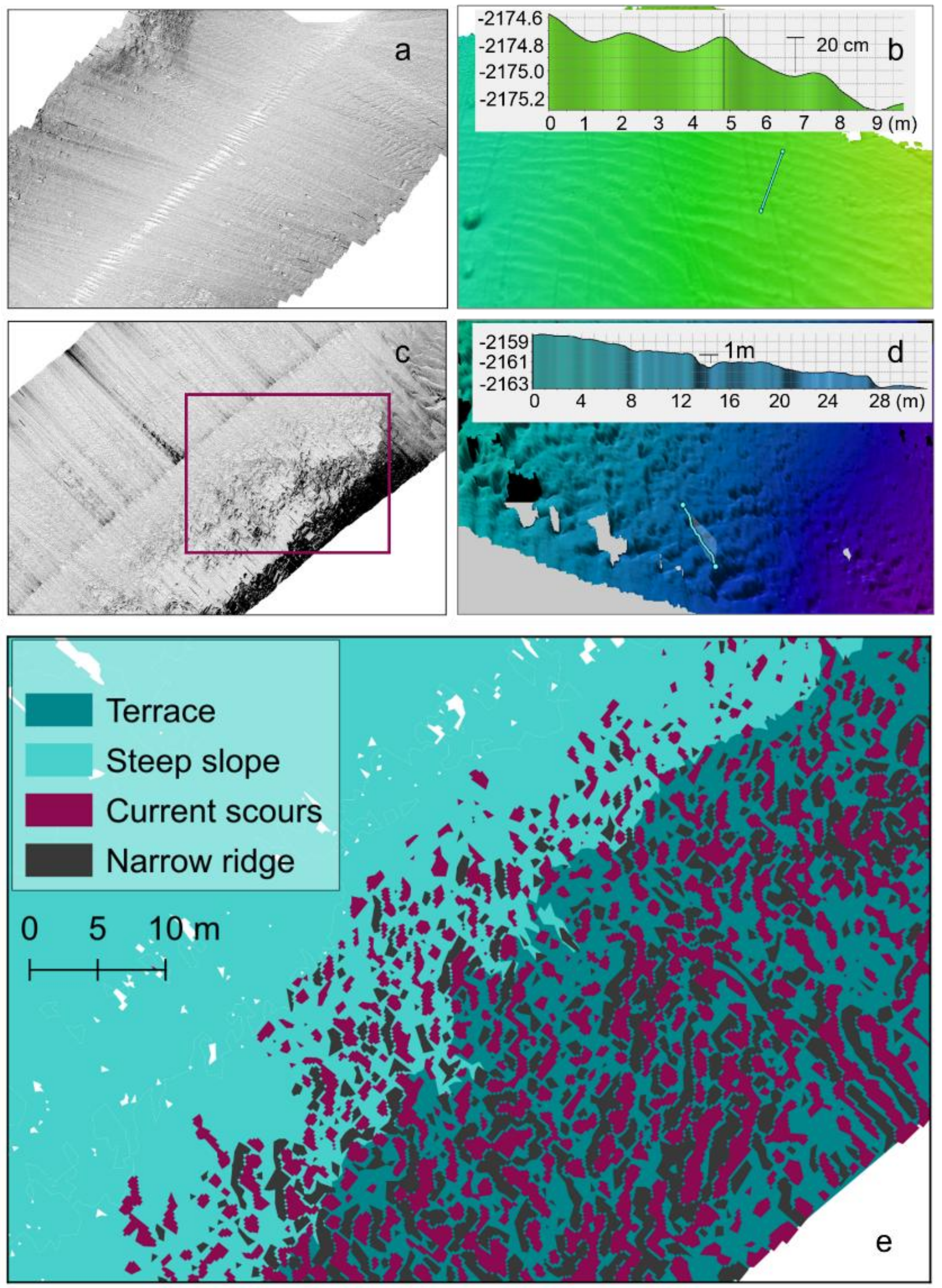


**Supplementary Figure S5: Landform terrain features.** a–b) Sand ripples illustrated by 3D bathymetry and sidescan. The insert in b) depicts ripple heights along a 9 m transect. c–d) Terraces illustrated by 3D bathymetry and sidescan. The insert in d) depicts terrace step heights along a 28 m transect. e) Zoomed-in view of the red box in (c), illustrating the topographic complexity on terraces with narrow ridges and current scours.

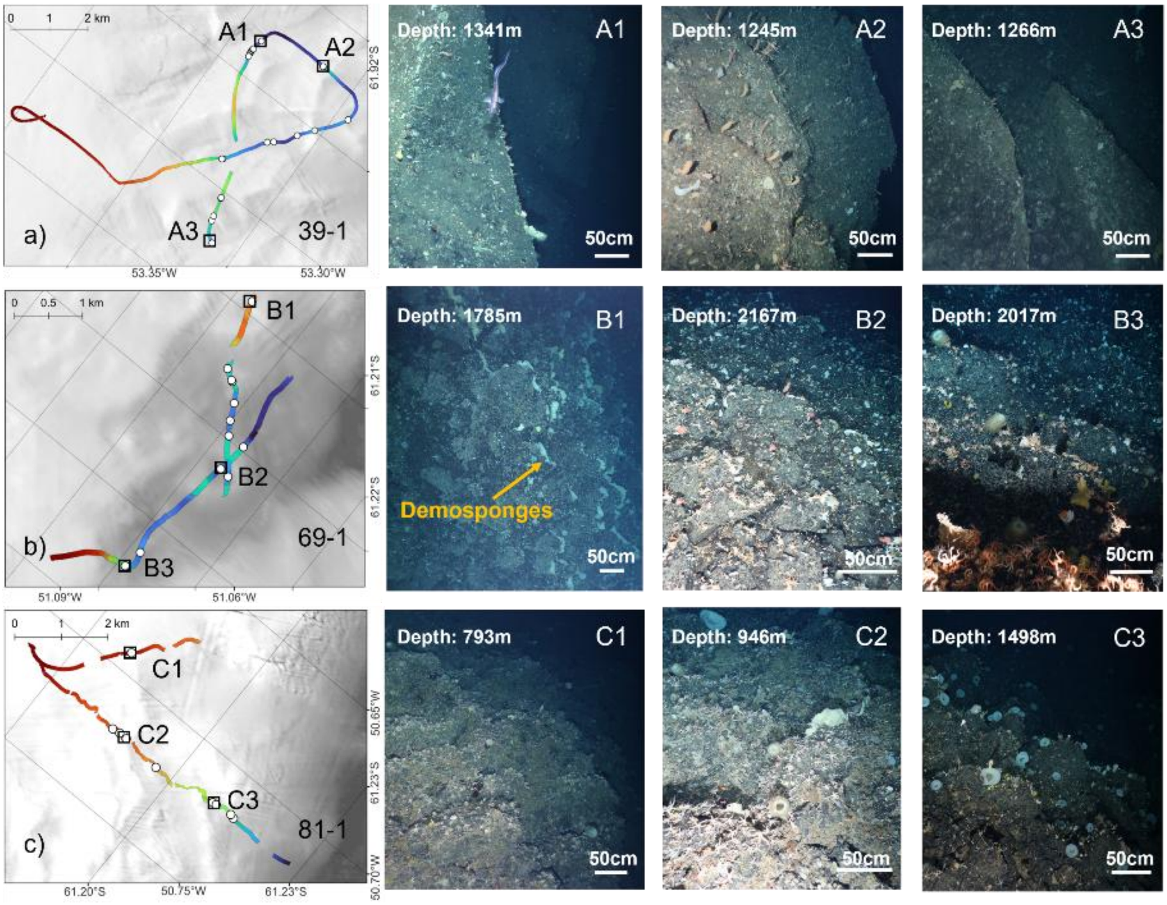


**Supplementary Figure S6: Terraces and epibenthos.** Locations of terraces on OFOBS dives 39-1, 69-1 and 81-1 (white dots in panels a-c) and example images of associated benthic communities (A1-C3). Arrows indicate demosponges on terrace edges.

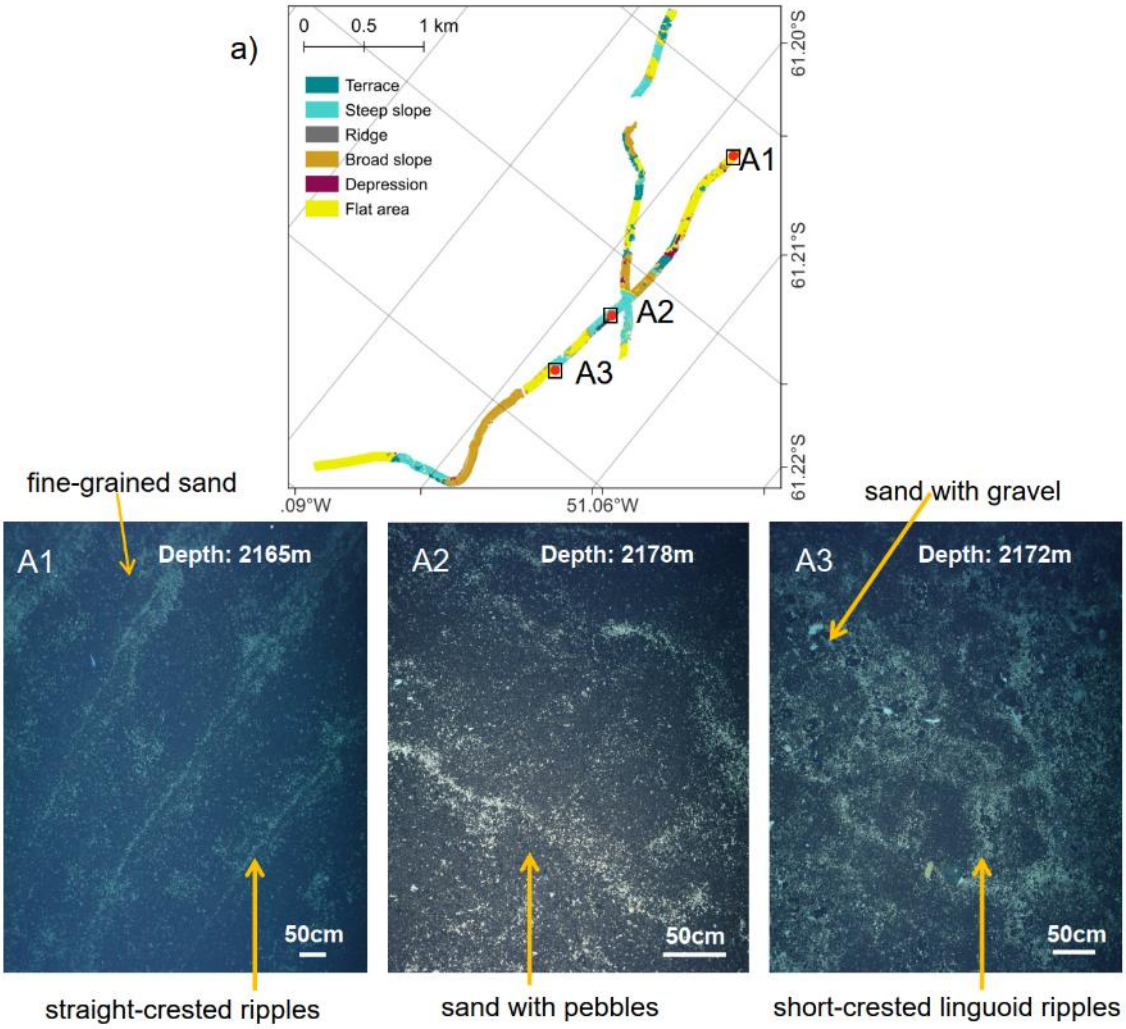


**Supplementary Figure S7: Sand ripples and epibenthos.** Locations of sand ripples in dive 69-1 (a), and example images of associated benthic communities (A1-A3). A1: fine-grained sand substrate with a straight-crested ripple pattern. A2: sandy substrate mixed with pebbles. A3: sandy substrate with gravel. Ripples in fine-grained sand indicate steady, slow bottom currents whereas linguoid ripples associated with gravel and pebble patches suggest increased flow velocities.

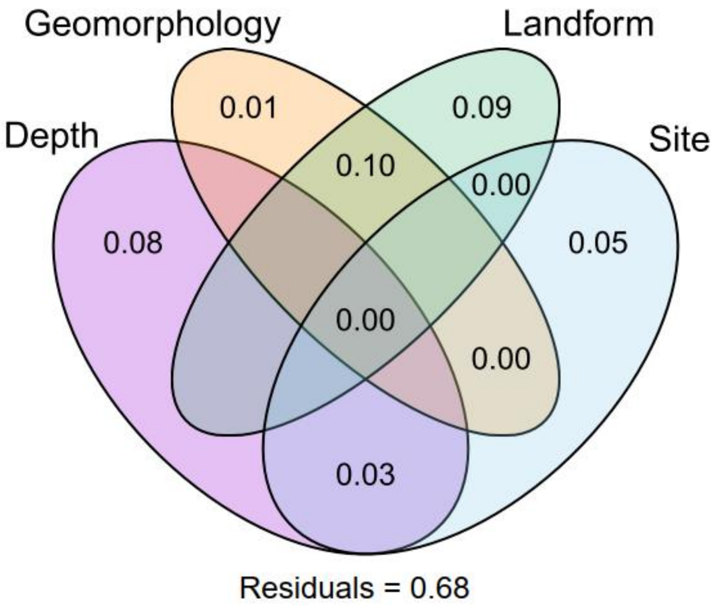


**Supplementary Figure S8: Variance partitioning of RDA predictors.** Venn diagram showing the relative contribution of variable groups on community discrimination between landforms in RDA.

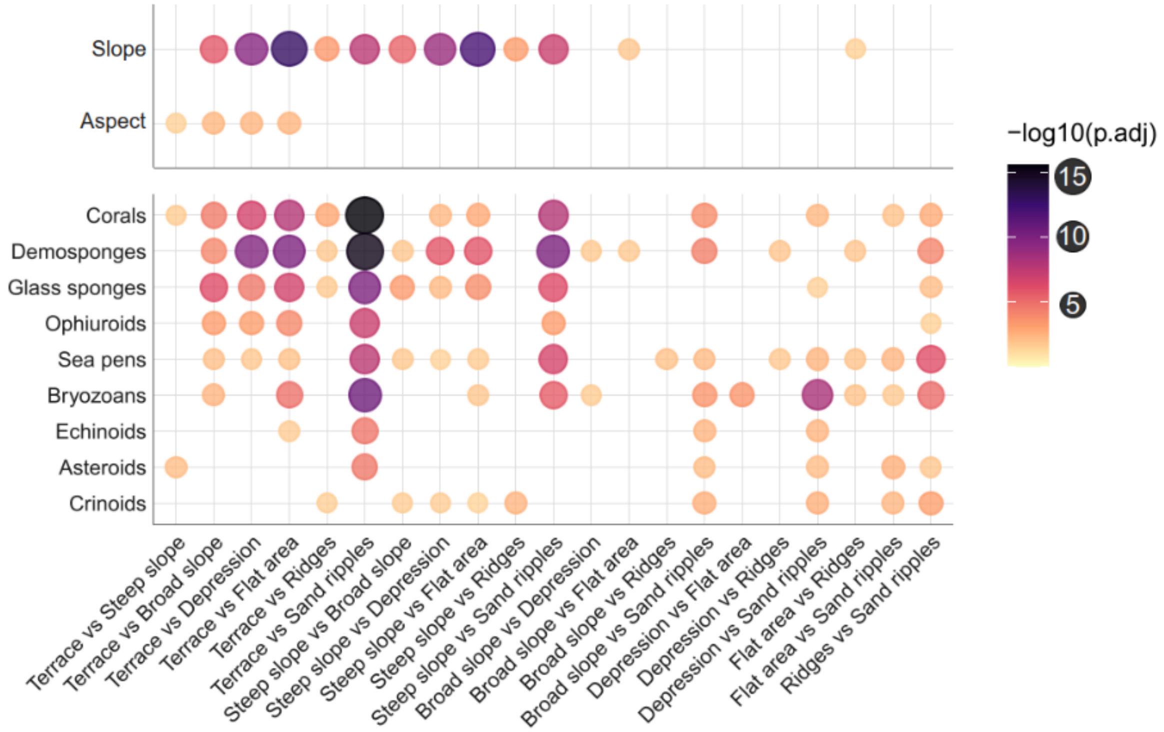


**Supplementary Figure S9:** Significantly differing terrain variables (top) and taxon abundances (bottom; Terraces and Steep slopes highlighted in green shades) between landforms.
